# The Genomics of Selfing in Maize (*Zea mays* ssp. *mays*): Catching Purging in the Act

**DOI:** 10.1101/594812

**Authors:** Kyria Roessler, Aline Muyle, Concepcion M. Diez, Garren R.J. Gaut, Alexandros Bousios, Michelle C. Stitzer, Danelle K. Seymour, John F. Doebley, Qingpo Liu, Brandon S. Gaut

## Abstract

In plants, self-fertilization is both an important reproductive strategy and a valuable genetic tool. In theory, selfing increases homozygosity at a rate of 0.50 per generation. Increased homozygosity can uncover recessive deleterious variants and lead to inbreeding depression, unless it is countered by the loss of these variants by genetic purging. Here we investigated the dynamics of purging on genomic scale by testing three predictions. The first was that heterozygous, putatively deleterious SNPs were preferentially lost from the genome during continued selfing. The second was that the loss of deleterious SNPs varied as a function of recombination rate, because recombination increases the efficacy of selection by uncoupling linked variants. Finally, we predicted that genome size (GS) decreases during selfing, due to the purging of deleterious transposable element (TE) insertions. We tested these three predictions by following GS and SNP variants in a series of selfed maize (*Zea mays* ssp. *mays*) lines over six generations. In these lines, putatively deleterious alleles were purged, and purging was more pronounced in highly recombining regions. Homozygosity increased more slowly than expected; instead of increasing by 50% each generation, it increased by 35% to 40%. Finally, three lines showed dramatic decreases in GS, losing an average of 398 Mb from their genomes over the short timeframe of our experiment. TEs were the principal component of loss, and GS loss was more likely for lineages that began with more TE and more chromosomal knob repeats. Overall, this study documented remarkable GS loss – as much DNA as three *Arabidopsis thaliana* genomes, on average - in only a few generations of selfing.

## INTRODUCTION

Darwin showed that the self-fertilization of plants leads to reduced vigor and fertility – i.e., inbreeding depression (Darwin, 1876). His work supported the hypothesis that self-fertilization is strongly disadvantageous and also provided a rationale for the prevalence of outcrossing in nature (Fisher, 1941, Morran et al., 2009). He did not, however, know the genetic basis of inbreeding depression. It is now thought to be caused by increased homozygosity, which inflates the genetic load by uncovering recessive deleterious alleles and/or by eliminating heterozygosity at loci with an overdominant advantage (Charlesworth and Willis, 2009, Hedrick and Garcia-Dorado, 2016). The increase of homozygosity – or, alternatively, the decrease of heterozygosity (*H*) - is expected to occur at a regular rate; in a selfed lineage, *H* is expected to be halved each generation. However, the actual rate of *H* decline is likely to be slowed by various factors, such as interference due to linkage (linked selection), epistatic interactions (Hedrick et al., 2016) and selective pressure to retain heterozygosity at overdominant and associative-overdominant loci (Byers and Waller, 1999, Schnable and Springer, 2013). These factors presumably contribute to the fact that inbred lines of maize and selfing *Caenorhabditis* species retain some heterozygosity, even after many generations of selfing (Barrière et al., 2009, McMullen et al., 2009, Rodgers-Melnick et al., 2015, Brandenburg et al., 2017).

One way to combat the increased load caused by inbreeding is the removal, or ‘purging’, of recessive deleterious alleles. When purging is effective, there may be no inbreeding depression (Crnokrak and Barrett, 2002). Purging is expected to occur rapidly when recessive alleles have lethal effects (Lande and Schemske, 1985, Charlesworth, 1990, Hedrick, 1994, Schultz and Willis, 1995) but should be less efficient for non-lethal recessives (Byers and Waller, 1999, Crow, 2008). The existence of purging is supported by experiments, theory and forward simulations (Charlesworth and Willis, 2009, Arunkumar et al., 2015, Liu et al., 2017), but it is expected to vary across species based on features like population history, mating system, and the distribution of fitness effects. Given this variation, one meta-analysis has concluded that purging is an “inconsistent force” in the evolution of inbreeding plant populations (Byers and Waller, 1999).

Recently, authors have argued that genomic data provide more precise insights into inbreeding effects than previous approaches (e.g. (Hedrick and Garcia-Dorado, 2016, Hedrick et al., 2016, Kardos et al., 2016)). Here we extend that argument to the phenomenon of purging, beginning with three simple predictions. The first is that selfed offspring will exhibit a bias against the retention of putatively deleterious SNP variants, because these SNPs become uncovered in a homozygous state. The second is that purging of SNP variants will be inconsistent across genomic regions, based on the amount of recombination. All else being equal, regions of high recombination should purge deleterious variants more efficaciously, because recombination reduces interference among selected sites (Hill and Robertson, 1966, Morran et al., 2010).

The third and final prediction is that purging will decrease genome size (GS). We make this prediction because GS correlates strongly with transposable element (TE) content (Tenaillon et al., 2010, 2011, Diez et al., 2014, Bilinski et al., 2018) and because plant TE insertions are thought to be predominantly deleterious (Wright et al., 2013). As a consequence, inbreeding should purge TE insertions by favoring the retention of haplotypes with fewer TEs. This may be especially true for TE insertions near genes, which are deleterious in part through their effects on gene expression (Hollister and Gaut, 2009, Hollister et al., 2011, Quadrana et al., 2016, Lee and Karpen, 2017). Consistent with these predictions, selfing species tend to have smaller genomes than outcrossers in both plants (Price, 1976, Govindaraju and Cullis, 1991, Wright et al., 2008) and animals (Fierst et al., 2015).

In this study, we take an ‘experimental evolution’ approach to investigate the dynamics of purging on a genome-wide scale. The experiment mimics an immediate transition to selfing, because it consists of 11 outcrossed maize parental lines that were self-fertilized for six or more generations. Given these selfed lineages, we gathered flow cytometric and whole genome resequencing data to address three sets of questions. First, does GS decrease rapidly in selfed lineages? If so, are TEs the primary component that is lost? Second, are putatively deleterious alleles purged more rapidly than putatively neutral alleles, and if so, does purging vary with recombination rate? Finally, does *H* decline at expected rates over time?

## RESULTS

### Plants, Phenotypes and Genome Sizes

The plant material came from a previous experiment in which 11 heterozygous maize landraces were self-fertilized to create homozygous lines (Wills et al., 2013). For each landrace, the experiment began with a single, outcrossed parent (P) of unknown genotype, and selfing was continued for ≥6 generations by single seed descent. For this study, we germinated seeds from intervening generations – i.e., from S1 to ≥S6. Each of our seeds was a sibling to the seed that was used to propagate the ensuing generation (Figure 1A). Following germination, we sowed 3 plants per line per generation. The plants did not flower under our growth conditions, but we measured growth rate and mortality (proxies for fitness) over a 45-day period. Growth rate and mortality varied among the eleven lines (p<0.001; **Figures S1 & S2**).

**Figure 1:**
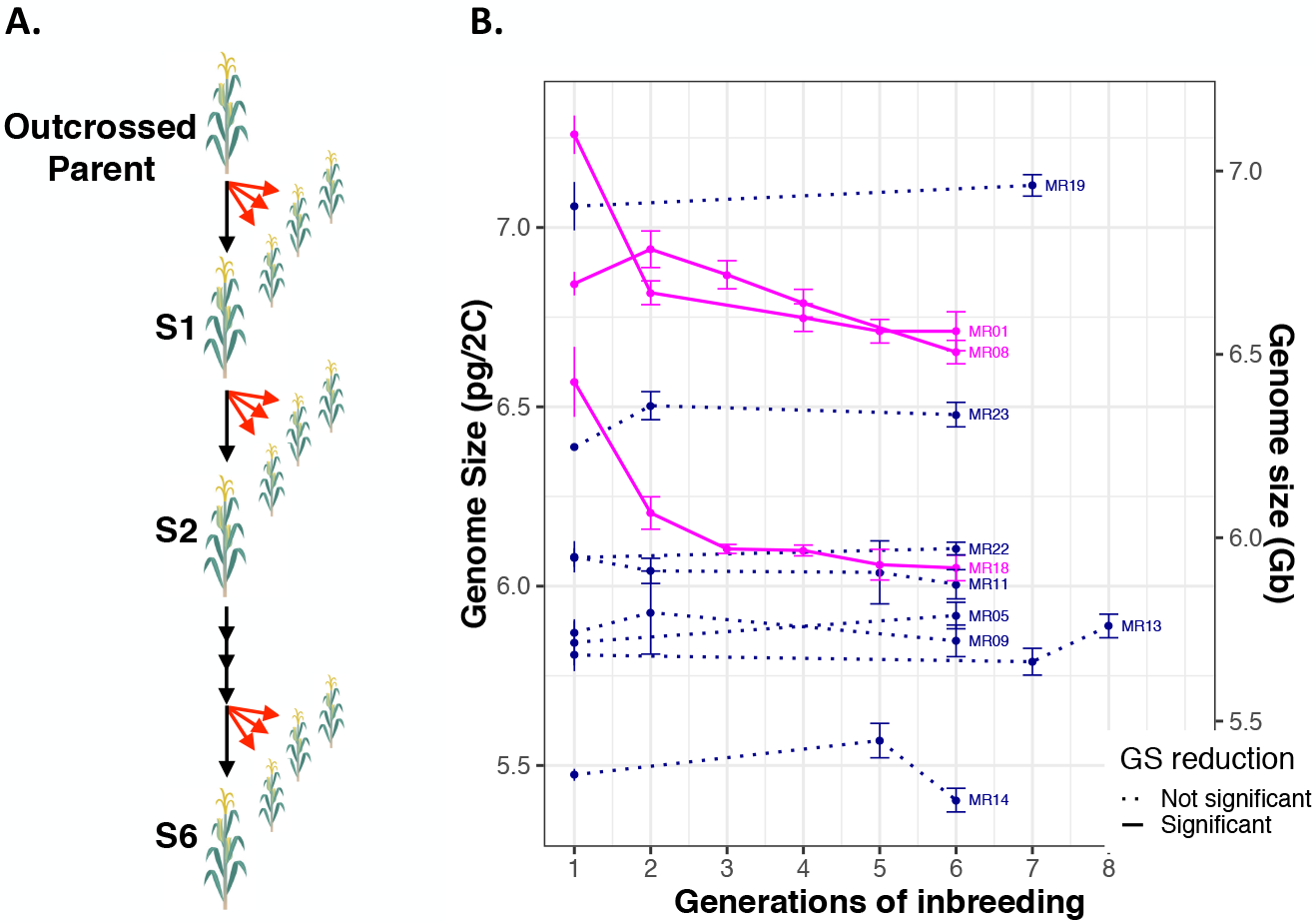
*A*) A schematic of the study design. An outcrossing parent was selfed to make the S1 generation and then subsequently selfed to S6 and higher. The selfed, single-seed descent lineages are represented by black arrows. Our study used sibling seed sampled from each generation, represented by red arrows. *B*) Estimates of genome size, in pictograms per 2C content, across generations of selfing. Each of the 11 lines is represented. Dark lines represent significant decreases of GS. Dotted lines did not have significant changes in GS. Mean and standard error are plotted. See Table S1 for sample sizes, Table S2 for raw values and Figure S3 for a detailed plot of the raw data per line.

To test for GS change, we gathered flow cytometry estimates for 96 plants and five B73 controls. Plant choice was restricted by mortality, but the 96 plants were chosen to represent a time series for each of the 11 lines, with > 1 plant per generation where possible (**Table S1**). We included three technical replicates per plant, for a total of 303 assays (**Table S2**). We then investigated our prediction of GS loss in two ways. First, we contrasted GS between the S1 generation and the latest (≥S4) generation with at least two siblings. By this measure, three lines (MR01, MR08 and MR18) exhibited significant decreases in GS (Wilcoxon rank-sum tests; *p* < 0.05), with no detectable GS shifts for the remaining eight lines (*p* > 0.5; **Table S3**). Second, we plotted flow cytometry data as a function of time, which included data from intermediate generations (Figures 1B **& S3**). The results again indicated that MR01, MR08 and MR18 exhibited significant decreases in GS and that the other lines had no detectable loss, based on linear and exponential model fits (**Table S4**). For MR01 and MR18, a model of exponential decay fit the data better than a linear model, suggesting that GS loss occurred more rapidly in the early generations.

We made three further observations based on flow cytometric data. First, GS loss occurred in three of the four lines with the largest S1 genomes (Figure 1B). These rankings were non-random by permutation test (*p* = 0.006), illustrating an increased tendency for lines with larger genomes to lose size. Second, because none of the lines exhibited a significant GS increase, the probability of GS loss was significantly higher than GS gain (*p*=0.04; two-sided binomial). Finally, we estimated the number of bases lost by each line, assuming a reference value of 5.64 pg/2C for maize B73 (Diez et al., 2013) and a conversion rate of 1pg = 978 Mb (Dolezel et al., 2003). Line MR01, for example, had an average GS estimate of 7.26 pg/2C in S1 and a corresponding average of 6.75 pg/2C in generation 4. The difference between generations was therefore 0.51 pg, which corresponds to a loss of 7.0%, or 499Mb. Similarly, lines MR08 and MR18 lost 2.8% (or 186Mb) and 7.9% (or 508 Mb) between generations 1 and 6.

### Genomic Components Correlate with GS Variation Across Samples

We predicted that purging leads to GS loss, which was true for 3 of 11 lines. We also predicted that loss would be dominated by TEs, but TEs are not the only potential genomic component that may contribute to rapid GS reduction. GS loss could also be attributed to: *i*) the loss of genes, *ii)* variation in rDNA copy number (Cullis, 2005, Long et al., 2013), *iii*) fluctuations in the number of chromosomal knob and CentC repeats (Jian et al., 2017, Bilinski et al., 2018); or *iv*) the loss of supernumerary B-chromosomes, which are small (Mroczek et al., 2006) but can be multicopy (Randolph, 1941) and vary among accessions (Yamakake, 1976).

To investigate the genomic regions responsible for GS change, we resequenced 33 plants that included data from S1 and ≥S5 for the three lines that exhibited GS loss (MR01, MR08 and MR18; the GSΔ group) and from three control lines (MR09, MR19 and MR22; the GS_con_ group) (**Table S1**). The data were mapped to the maize B73 AGPv4 genome with four annotated genomic components -- genes, rDNA, TEs, knob-specific repeats – and B chromosome repeats (see **Methods**). Total read counts varied among individuals; hence comparison across individuals and generations required normalization. Similar to previous papers (Diez et al., 2014, Bilinski et al., 2018), we normalized across libraries based on the number of read counts to genes, but in this case we focused on single copy BUSCO orthologs (Simão et al., 2015)(see Methods). Our reasoning was that BUSCO genes were unlikely to contribute to short-term GS change, because they are conserved across the kingdom Plantae. Simulations demonstrated that this normalization approach leads to accurate inferences of relative read counts in genomic components (like TEs) that may vary across generations (**Figure S4**), even with low (2x) coverage.

Given normalized read count data, we examined the relationship between GS (as measured by flow cytometry) and sequence counts across the entire sample of 33 plants. Regressing each component separately, there was no significant relationship to GS for genic content (*r*^2^=−0.027, p=0.63) or B-chromosome content (*r*^2^=−0.015, *p* = 0.45). There was borderline significance for rDNA (*r*^2^=0.079, *p*=0.07), but strongly positive relationships between GS and both knob repeat content (*r*^2^=0.662, *p*=4.5×10^−8^) and TE content (*r*^2^=0.901 p< 10^−15^**; Figure S5**). When all of the components were combined into a single linear model, only TE counts remained significant (linear model t-value= 9.18, *p*= 2.55 ×10^−09^), but knobs were again significant after TE counts were removed from the model (linear model t-value= 5.78, *p*= 5.02 ×10^−06^). Hence, GS correlates most strongly with TE content but there is a hint that knobs also contribute to GS variation.

### Genomic Components that Contribute to Temporal Loss

TEs and knobs contribute to GS variation, but which among the five components varied over time and contributed to GS change? To address this question, we applied ANOVA to read count data from each of the five genomic components separately. The ANOVA tested for significant differences between *groups* (GSΔ vs. GS_con_), among *landraces* (e.g., MR01 to MR22), and between *generations* (e.g., S1 to S6). It also tested for *group*generation* and *landrace*generation* interactions. We were particularly interested in *group*generation* interactions, because they identify components that differentiate the GSΔ vs. GS_con_ groups over time.

We applied ANOVA to each of the six genomic components separately (Tables 1 **& S5**) and plotted normalized counts for groups (Figure 2) and landraces (**Figure S6**). Focusing first on genes, the ANOVA had no significant terms (*p* > 0.05; all p-values FDR corrected for all of the tests in Table 1). The lack of significance was reflected in plots of read counts, because there were only moderate differences between groups and among landraces, without a consistent trend over time. For rDNA, the ANOVA detected differences among landraces (F-value= 5.28, *p*=0.004), with 41% of the variance explained (VE) but with no other significant terms. By comparing GS estimates to read counts (**see Methods**), we estimated the average number of Mb’s attributable to rDNA repeats in each line and each generation. No line had > 8Mb of estimated rDNA, and the temporal difference between S1 and S6 was < 0.7 Mb for most lines (**Table S6**). A third component was B-chromosomes. Only one line (M18) had substantial hits to B-chromosome repeats, representing an average of 10.7 Mb of DNA content across S1 individuals. By S6, counts were at background levels, indicating the loss of B-chromosomes. Given these patterns, the ANOVA detected significant *landrace* (F-value=5.90, *p* =0.021) and *landrace*generation* terms (F-value=4.85, *p* < 0.022), but no group effects.

**Table 1:** Estimates of the variance components based on ANOVA applied to read count data. Each of the five genomic components (TEs, genes, knob-repeats, B chromosome specific repeats and rDNA) was tested individually.

|  | **Group** | **landrace** | **generation** | **Group X gen** | **Line X gen** |
| --- | --- | --- | --- | --- | --- |
| **TEs** | 14.72 *** | 70.65*** | 2.82* | 5.41* | 0.65 |
| **Genes** | 1.46 | 21.49 | 4.85 | 0.017 | 18.51 |
| **Knobs** | 35.60 *** | 56.21*** | 0.44 | 0.50 | 2.54 |
| **bChr** | 7.02 | 25.80* | 7.27 | 6.76 | 25.53* |
| **rDNA** | 2.00 | 40.49* | 2.27 | 1.53 | 13.47 |

<sup>1</sup> Statistical significance is indicated by \* < 0.05; 0.05 > \*\* >0.001, \*\*\*<0.001. P-values were FDR corrected based on all tests in the Table.

**Figure 2:**
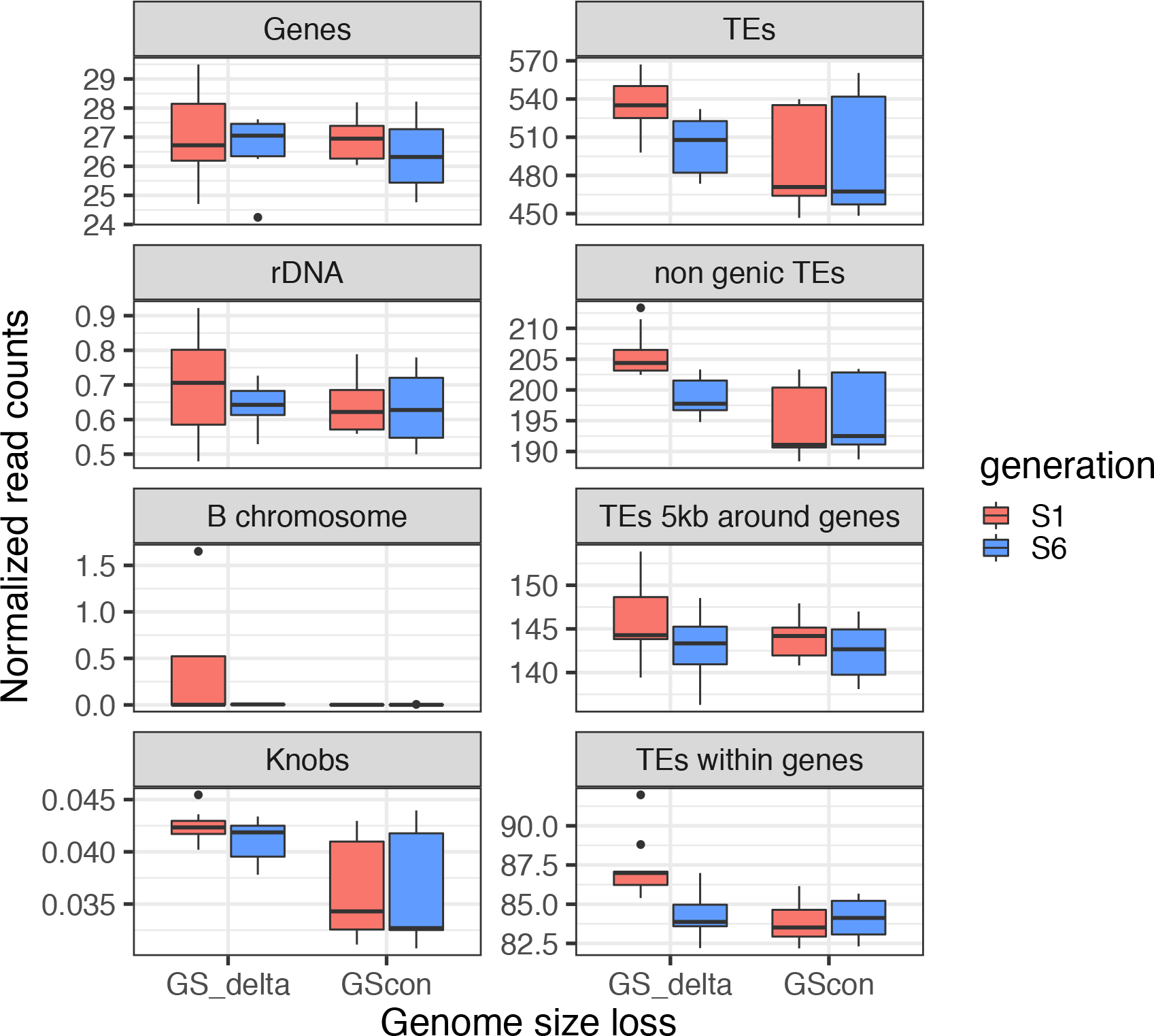
Various components of the genome compared between the GS change group (GSΔ) and the GS constant (GS_con_) groups and between S1 and S6. Sample sizes are shown in Table S1, significance values are provided in Table **S5**, and Figure **S6** reports this information for each of the lines separately. The boxplot shows the median, lower and upper quartiles. The whiskers extend to the largest or lowest value no further than 1.5 * IQR from the hinge (where IQR is the inter-quartile range, or distance between the first and third quartiles). Outliers are plotted as dots above the whiskers.

We next turned to the two genomic components that correlated strongly with GS across the entire dataset: TE counts and knob repeats. TE counts exhibited significant terms across *groups* (F-value=53.94, *p*=2.38×10^−07^; 14% VE), *landraces* (F-value=64.71, *p*=2.91×10^−11^; 70% VE), *generations* (F-value=10.35, *p*=0.018; 2.8% VE) and *group*generation* interactions (F-value=19.84, p=0.0013; 5.4% VE)(Table 1). The plots of TE counts were consistent with these statistical results, because they show that: *i*) the GSΔ group had higher overall TE counts than the GS_con_ group; *ii*) landraces within GSΔ exhibited marked reductions in TE counts from generation S1 to S6, but *iii*) landraces within GS_con_ did not (Figure 2). By equating GS to read counts, we estimated that the Mb loss due to TEs was 481 Mb for MR01, 199 Mb for MR08 and 465 Mb for MR18, representing >90% of the estimated shift in GS over time for each line. In contrast, the GS_con_ lines exhibited temporal TE changes of ~10Mb each (**Table S6**).

Finally, knob counts differed between groups (F-value=158.99, *p*=2.91×10^−10^; 56% VE) and among landraces (F-value=62.75, *p*=2.91×10^−10^; 35% VE), with the GSΔ group having generally higher counts. However, knob counts did not exhibit significant interaction terms or variation between generations, which was surprising given the correlation between knob counts and GS across all samples (**Figure S5**). We thus investigated the possibility that the lack of significance reflected reference bias by repeating analyses with the W22 reference (Springer et al., 2018). The results largely corroborated the B73 results but did produce a significant *group*generation* interaction for knobs (F-value=10.88, *p*=0.0128) (**Table S7**). Based on the W22 reference, the average Mb loss over generations due to knobs was 136 Mb in MR01, 59.4 Mb in MR08 and 77.0 in MR189, but TEs explained more temporal variation in every case (341 Mb, 130Mb, and 413 Mb, respectively; **Table S8**).

### TE Locations, Types and Mechanism

We predicted that GS loss could reflect purging of TEs near genes due to their deleterious effects on gene expression (Hollister and Gaut, 2009, Lee and Karpen, 2017). To address this prediction, we separated TEs from B73 into three bins: non-genic TEs, which mapped to TEs > 5kb away from genes; near-genic TEs that were within 5kb of a gene; and the subset of near-genic genes that overlapped with annotated genes – i.e., they fell within introns or UTRs. Both non-genic and overlapping TEs exhibited significant *group*generation* interactions (F-values=18.46 and 13.97, p≤0.001 all p-values FDR corrected; **Table S9**), explaining 9.1% and 12.4% of the total variance for non-genic and overlapping TEs, respectively. Despite our prediction, none of the ANOVA terms were significant for TEs near (< 5kb) genes, but three components were borderline significant (F-value=3.34, p < 0.10; **Table S10**), including both the *generation* and *landrace*generation* terms. Interestingly, the latter reflects the fact that five of the six lines lost near-genic TEs through the course of the experiment (**Figure S6**), suggesting that the loss of TEs near genes was a general phenomenon across all lines. We repeated these analyses for the W22 reference, and we found that all three TE locations exhibited *group*generation* effects (**Table S10 & Figure S7**). Overall, then, these data suggest that TEs were lost throughout the genome, but it is unclear whether near-genic TEs were lost across all lines or only from the GSΔ group.

We also investigated potential biases by TE order, focusing on six TE types in the B73 reference: Helitrons, Long Terminal Repeat (LTR) retrotransposons, solo LTRs, Terminal Inverted Repeats (TIRs), SINEs and LINEs. All but solo LTRs exhibited significant variation between the GSΔ and GS_con_ groups (F-value > 39.10, *p*<1.2×10^−5^). Four of the six also exhibited a significant *group*generation* interaction, which explained > 5% of the variance for LTRs, solo LTRs and Helitrons (**Figure S8 & Table S11**). Thus, GS loss encompassed an array of TE types.

Finally, we addressed a question related to a potential mechanism of TE loss. In some plant species, TE loss is driven by unequal recombination between LTR elements (Devos et al., 2002). These recombination events are expected to increase the ratio of solo LTR elements to intact LTR elements. If this mechanism operated during our experiment, the ratio of reads mapping to LTRs vs. the internal regions of elements should increase over time, especially in the GSD lines. To test this idea, we independently annotated 22,530 full-length LTR elements of the Sirevirus genus, based on the B73 reference. We focused on Sireviruses for three reasons: *i*) they represent a substantial proportion (~20%) of the maize genome (Bousios et al., 2012), *ii*) they can be accurately annotated based on numerous internal features, including the boundary between LTRs and internal regions (Darzentas et al., 2010), and *iii*) they provide a set of LTR elements that were annotated independently of the existing B73 v4 genome annotation. We found that both solo and intact Sireviruses exhibited losses over time in the GSΔ group (**Table S12**; **Figure S9**), which is consistent with our LTR analyses based on the v4 annotations. However, the ratio of mapping to LTRs vs. internal regions did not exhibit an obviously increasing trend through time or a significant *group*generation* effect (F-value=0.27, *p*=0.73), as would be predicted if TE loss were driven by numerous unequal recombination events.

### The fate of deleterious variants

We now turn to a second prediction about purging: Over time, there should be a bias against the retention of deleterious SNP variants. We tested this prediction by first calling SNPs for each of the six lines from the GSΔ and GS_con_ groups and then by focusing only on bi-allelic SNPs that were inferred to be heterozygous (*H* = 1) in the resynthesized parent (see **Methods**). For each of these heterozygous sites, we predicted derived deleterious variants using SIFT (Ng and Henikoff, 2003) and noted the fate of variants in four functional classes (non-coding, synonymous, tolerated nonsynonymous and putatively deleterious non-synonymous variants). In total, we examined 1,914,845 SNPs across the six lines (**Table S13**).

As a signal of purging, we expected deleterious, derived SNP variants to exhibit biased rates of loss over time. To characterize this potential bias, we identified derived alleles by comparison to *Sorghum* outgroup and estimated the proportion of derived allele (*P*_*d*_) across sites. We expected *P*_*d*_ to be 50% in the parent and to remain 50% in the absence of perturbing factors like selection. To test this prediction, we combined results across the six lines and plotted *P*_*d*_ for each functional class in S1 and S6 (Figure 3A). In S1, for example, average *P*_*d*_ estimates for non-coding and synonymous sites were below 0.5, potentially reflecting biases in ancestral inference and/or selection against a subset of these putatively ‘neutral’ derived alleles between Parents and S1. Consistent with the latter interpretation, *P*_*d*_ declined from S1 to S6 for both site classes (linear model contrast Z-value=14.92, *p* < 0.001).

**Figure 3:**
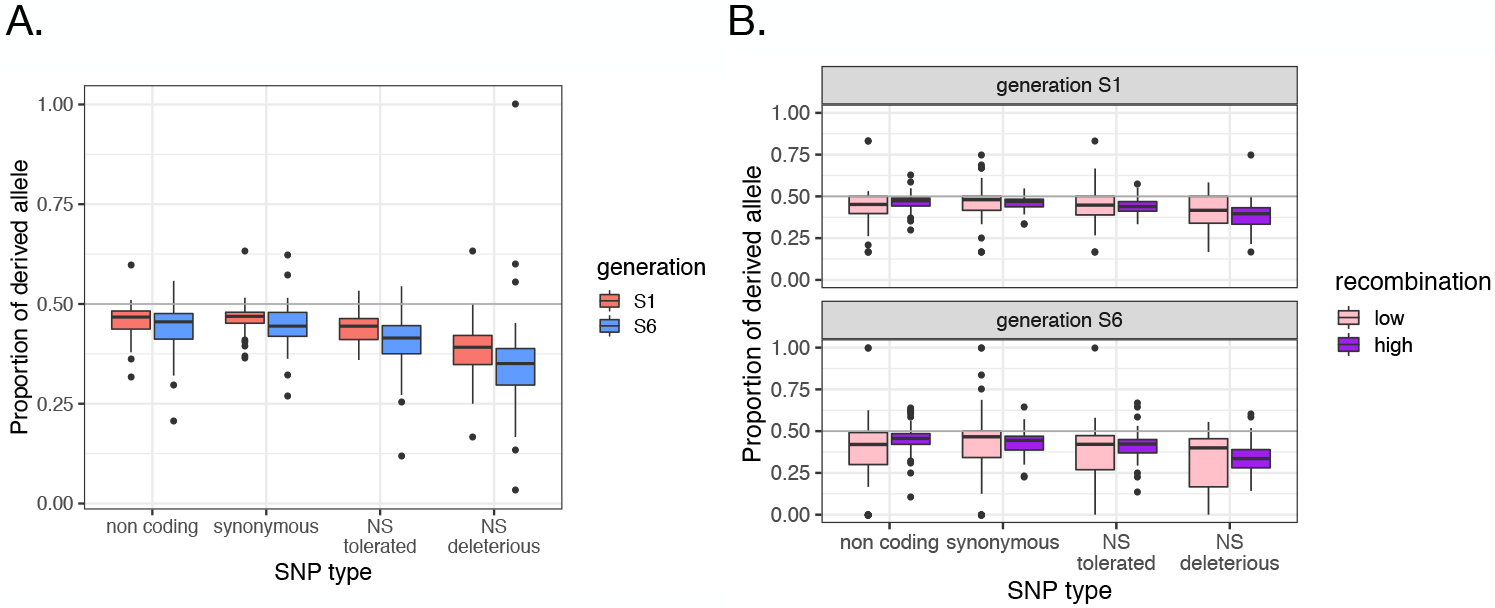
**A)** The proportion of the derived allele for the four mutational classes predicted by SIFT – i.e., non-coding, synonymous, non-synonymous tolerated and non-synonymous deleterious. The graph reports the proportion for generations S1 and S6 across six lines (MR01, MR08, MR09, MR18, MR19 and MR22). P_D_ was averaged across individuals for each chromosome and line separately (n=60 for each bar of the plot, n=480 in total). **B)** As in panel A, except the genome was separated into high and low recombination quartiles of the genome, illustrating that purging occurs more rapidly in high recombination regions. As in A), n=60 for each bar of the plot. See Figure 2 legend for values of the boxplot.

Importantly, these effects were greatly amplified for nonsynonymous mutations (Figure 3A). For example, putatively deleterious, derived nonsynonymous SNPs had a *P*_*d*_ of 0.384 in S1, representing a significant decrease relative to that of synonymous and non-coding variants (linear model contrast Z-value=44.89, p< 0.001; **Table S14)**. Between S1 and S6, *P*_*d*_ fell even further, from an average of 0.384 to 0.334 (linear model contrast Z-value=20.83, p<0.001). Overall, putatively deleterious SNPs demonstrated accelerated rates of loss over time relative to other variant classes.

Recombination is expected to mediate the effects of selection, because it uncouples interference between linked variants. Therefore, deleterious variants should be purged more rapidly in regions of high recombination. To explore this prediction, we contrasted genomic regions that encompass the highest and lowest quartiles of recombination rates, as defined by cross-over events (*r*) (Rodgers-Melnick et al., 2015). The results showed the expected pattern: in S6, *P*_*d*_ was lower in high compared to low recombining regions for both classes of nonsynonymous variants (tolerated: Z-value=−3.37, *p*=0.006; deleterious: Z-value=−4.95, *p* = 5.98×10^−6^ based on linear model contrasts; **Table S15**). Recombination did not have an effect on *P*_*d*_ for nonsynonymous SNPs in S1, consistent with the fact that time is required for recombination to break down linkage between loci.

### Declining Heterozygosity

Finally, we measured a phenomenon of empirical interest, which is the rate of loss of heterozygosity over generations. To do so, we took advantage of the fact that SNPs inferred to be heterozygous in the parental generation can be in only one of two states within S1 and S6: heterozygous or homozygous. Moreover, these two states are expected to fall into blocks, with the transition between blocks defined by recombination events. To identify these blocks, we examined windows of 100 SNPs in size, focusing on genic SNPs, and used a Bayesian clustering method to assign windows as either heterozygous or homozygous for each individual (see **Methods**). The proportion of heterozygous SNPs across the genome (*H*_*b*_) can be compared directly to the null expectation that *H* = 0.50 in S1 and 0.015 in S6.

We applied this approach successfully to the two lines with highest coverage (MR09 and MR22) (Figure 4) and offer five observations about heterozygosity. First, *H*_*b*_ exceeded 60% in both MR09 (65.7%) and MR22 (63.7%) for generation S1, representing a significant deviation from the null expectation (one-sided Wilcoxon test p=0.0019 and p=0.019 respectively). Second, *H*_*b*_ significantly exceeded the expected value of 1.5% in S6, at 14.2% for MR22 and 4.8% for MR09 (one-sided Wilcoxon test p=0.00098 and p=0.019 respectively). Third, for reasons that are not immediately apparent, the difference between the two lines in S6 was also significant (one-sided Wilcoxon test *p* = 0.00036). Fourth, heterozygous blocks had a significantly higher proportion of nonsynonymous SNPs (7.19%) compared to homozygous blocks (6.14%, one-sided chi-square = 27.72, *p* = 1.4×10^−7^). Finally, heterozygosity was also related to recombination, because heterozygosity and *r* were modestly but significantly correlated across windows in S6 (linear regression adjusted *r*^2^ = 0.016; p = 1.5×10^−4^).

**Figure 4:**
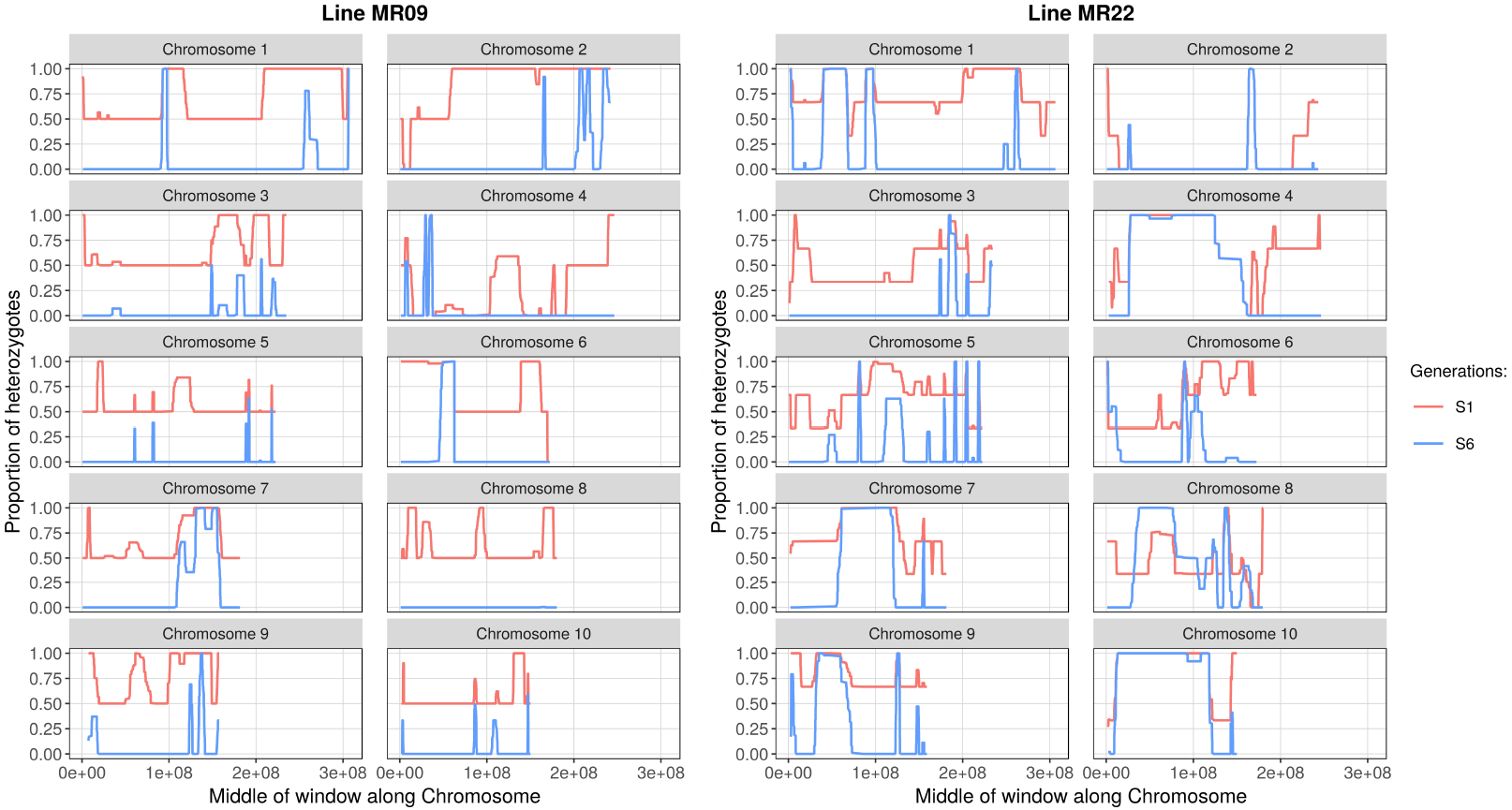
Inference of heterozygous and homozygous genomic regions, based on SNPs inferred to be heterozygous in the Parent. The figure shows each of the ten chromosomes for two lines (MR22 and MR19). Heterozygosity was averaged across individuals for each line and generation separately. For each chromosome, the x-axis represents length along the chromosome and the y-axis is the proportion of heterozygous sites within 100 SNP sliding windows. Red and blue lines represent the S1 and S6 generations. Both lines have more regions of heterozygosity than expected (see text for statistics). Sample sizes are shown in Table S1 (n=2 or 3 depending on the line and generation).

## DISCUSSION

Self-fertilization is an important reproductive strategy in plants (Charlesworth and Wright, 2001), and it is also a widely applied tool for plant genetics and plant breeding. In this study, we took an experimental approach to assess the genomic effects of selfing, with a focus on the dynamics of purging. Previous studies have investigated the effects of selfing by, for example, contrasting selfing and outcrossing plants in flowering phenology, population structure, genomic diversity (Wright et al., 2013) and evolutionary fate (Takebayashi and Morrell, 2001). Yet, most of these effects likely accrue after, not during, the transition to selfing. A smaller number of studies have found evidence of purging by comparing inbreeding depression between naturally inbreeding and naturally outcrossing species (Crnokrak and Barrett, 2002, Weller et al., 2005, Charlesworth and Willis, 2009). In contrast, the immediate genomic effects of purging have gone largely undocumented.

### Rapid genome flux

Our experiment documents rapid GS loss in three of 11 selfed lineages (Figure 1). These observations add to a growing consensus that GS can change rapidly in plant species. Other examples include GS changes in flax over a single generation (Cullis, 2005), GS shifts on experimental time-scales in *Festuca* (Smarda et al., 2010), and GS reductions in maize after six generations of selection for early flowering (Rayburn et al., 1994). However, the magnitude of our observed GS losses is unprecedented. Based on flow cytometry estimates, the three lines lost ~6% of their genome, or 398 Mb, on average from S1 to S6. To put these changes in context, the GS of two fully-sequence maize inbred lines (Mo17 and B73) differ by only ~25Mb (Sun et al., 2018).

Following precedence (Tenaillon et al., 2011, Chia et al., 2012, Diez et al., 2014, Bilinski et al., 2018, Lyu et al., 2018), we used read counts to infer the size of genomic components, focusing on genes, TEs, knob repeats, rDNA and B-chromosomes. Among these five, it is clear that TEs are the major source of loss, which is not too surprising given that DNA derived from TEs constitute >85% of the maize genome (Schnable et al., 2009) and that previous studies have shown TEs contribute to plant GS variation (Tenaillon et al., 2011, Chia et al., 2012, Diez et al., 2014, Bilinski et al., 2018, Lyu et al., 2018). However, GS shifts are not always caused by TE content. In flax and *Arabidopsis thaliana*, GS shifts are fueled primarily by variation in rDNA repeats (Cullis, 2005, Long et al., 2013), and GS differences between selfing and outcrossing *Caenorhabditis* species are roughly equally apportioned among genes and TEs (Fierst et al., 2015, Yin et al., 2018).

Given that TEs are the major source of GS loss, we examined loss according to both TE type and location. Read count data indicate that loss occurred within the GSΔ group for all of the six TE orders we tested (**Figure S8** and **Table S6**). This finding – i.e., that TE loss is general and not limited to specific families - mirrors previous studies that have compared TE content among *Zea* genomes (Tenaillon et al., 2011, Diez et al., 2014). For example, Tenaillon et al. (2011) compared genome content between *Zea luxurians* and maize B73, two taxa that diverged ~140,000 generations ago (Ross-Ibarra et al., 2009). They estimated that 70% of the GS difference between species was due to TEs, but that the relative abundance of TE families was conserved between species.

We also predicted that GS loss should be especially evident for TEs that are near genes, because they are known to have deleterious effects on gene expression and genome function (Hollister and Gaut, 2009, Lee and Karpen, 2017). The results varied somewhat depending on the reference. With B73, the *line*generation* effect for near-genic TEs was borderline significant (*p* = 0.058), because five of the six resequenced lines lost these TEs over time, irrespective of their inclusion in the GSΔ or GS_con_ group (**Figure S5**). This result implies that the loss of near-genic TEs may be a general property of selfing. However, the W22 results do not fully support this claim, because they suggest that the pattern of loss in near-genic TEs varied between groups. Given these results, we cannot yet conclude that the loss of near-genic TEs is a general outcome of selfing. We advocate further investigation of this issue that also considers the fact that TE families vary in both their tendency to insert near genes and their epigenetic profiles.

In this context, it is important to emphasize the limitations of the read-count approach for estimating genomic components. The approach is better suited for broad-scale inferences about genome content than for inferences about the fate of specific genes, TE insertions or chromosomal regions. Here our inferences about location are based on the reference genome and may not accurately reflect the genome of our sample. We investigated reference biases by applying our read-count approach to two references (B73 and W22). With either reference, there was little evidence that genes, rDNA and B-chromosomes contributed substantively to GS loss, but the magnitude of the TE and knob effects did vary by reference. With B73, TEs explained > 90% of loss from S1 to S6 and as much as 481 Mb. With W22, the estimated TE loss was more modest, explaining ~75% of GS shift on average, with the remainder of loss assigned to knob repeats. The difference in results probably reflects annotation and assembly differences between references, because we disregarded counts from regions where annotated features overlapped. In B73, TEs often overlapped with putative knob regions, but overlaps occurred less frequently in W22. Our results therefore contain a cautionary tale about annotation biases, but we also suspect that the implication of knobs as a component of GS loss is reasonable, given our own (Figure S5) and previous evidence that knobs contribute to maize GS variation (Chia et al., 2012, Bilinski et al., 2018). Importantly, the total Mb loss explained by TEs and knobs was consistent, regardless of the reference.

Altogether, our results support the claim that GS loss is a common outcome of selfing, which is based on the fact that genomes are smaller in selfers versus their outcrossing sister taxa in *Caenorhabditis* (Fierst et al., 2015, Yin et al., 2018) and across plant taxa (Wright et al., 2008). And yet, if GS loss is common during selfing, one must wonder why 8 of our 11 lines exhibited no detectable loss. The lack of loss is probably not a question of statistical power, because five lines were estimated to have slightly *larger* GS, on average, in S6 relative to S1 (Figure 1). Here our lack of the parental genome could be misleading, because our experimental design could not monitor loss from the Parent to S1. Furthermore, we expect the greatest loss to be within the early generations, since two of the three GSΔ lines lost GS exponentially over time. We can nonetheless glean some predictive insights by contrasting data between GSΔ and GS_con_ groups. For example, neither group exhibited particularly low growth rates or high mortality (**Figures S1 & S2**), so GS loss did not obviously relate to these fitness proxies. The three lines with GS loss did have larger S1 genomes (Figure 1B), with significantly more TEs and knobs than the GS_con_ group (Figure 2; Table 1). To a first approximation, genomes with high TE and knob content are more prone to loss.

### Heterozygosity, recombination and the fate of deleterious SNPs

Several previous studies have shown that *H* declines at lower rates than expected under selfing (Hedrick et al., 2016), (Barrière et al., 2009). In S1 eucalyptus trees, for example, average *H* was 65.5%, compared to the expectation of 50% (Hedrick et al., 2016). We also find elevated heterozygosity in our lines. In S1, for example, *H*_*b*_ was ~ 65% for MR09 and MR22 (Figure 4). By S6, both lines retained significantly more heterozygosity than the expected value of 1.5%. Observed values of *H*_*b*_ in S6 imply that, assuming constancy across generations, the rate of heterozygosity retention was 0.60 per generation (= e^(log(0.048)/6)^) for MR09 and 0.72 per generation (= e^(log(0.142)/6)^) for MR22.

What can account for this remarkable retention of heterozygosity? One prosaic explanation is errors in heterozygosity assignments. While this is possible with our low coverage data, features of our design and analyses suggest that genotyping errors are not driving the results. First, if heterozygotes are miscalled as homozygotes in S1 due to low coverage, then they are simply ignored and do not affect the results. Second, we have taken advantage of the unique features of our experiment to infer heterozygosity in blocks; our results do not rely on heterozygosity calls at single SNPs. Third, we found no correlation in the location of heterozygous windows between MR09 and MR22 (r_2_ = 0.03846, *p*= 0.5871), suggesting that underlying genomic features (e.g., sets of paralogs that can cause SNP miscalls (Brandenburg et al., 2017)) did not consistently inflated heterozygosity between lines. Finally, we have made conservative assumptions about the categorical assignment of blocks (see **Methods**).

We are left, then, with the need to find biological explanations for slower-than expected rates of heterozygosity decline. Such explanations usually invoke either overdominance or associative overdominance (Ohta, 1971), with the prevailing view that associative overdominance is the prevailing force maintaining heterozygosity in selfed lineages (Springer and Stupar, 2007, Barrière et al., 2009, Charlesworth and Willis, 2009, Hedrick et al., 2016). Importantly, associative overdominance should hold for deleterious alleles with small phenotypic effects (Thornton et al., 2013).

If higher-than-expected levels of heterozygosity are caused in part by linkage to deleterious variants, then heterozygosity should be higher in regions of low recombination, where selection against deleterious variants is inefficient because loci are coupled. Consistent with this prediction, heterozygosity is elevated in regions of low recombination in the maize Nested Association Mapping (NAM) population (Gore et al., 2009, McMullen et al., 2009). In contrast, a study of deleterious variants in 247 inbred maize lines found little correlation between recombination rates and the proportion of deleterious SNPs, suggesting low recombination regions have enough recombination to purge deleterious variants over longer time periods (Mezmouk and Ross-Ibarra, 2014). Here, over the short-term timescale of our experiment, we find more nuanced and complex relationships with recombination. Overall, heterozygosity is lower in regions of low recombination, which probably reflects background selection (Charlesworth et al., 1993). In our experiment, we suspect that selection has acted against highly deleterious, recessive variants, potentially removing heterozygosity along large swaths of the genome. It is worth emphasizing that the correlation between heterozygosity and recombination is not due to inherently lower genetic diversity in low recombination regions of the parent, because our metrics focus only on the fate of inferred heterozygous sites in the parent, not on diversity per se.

Another feature of recombination is that it has the capacity to uncouple linked variants, making selection more efficacious. We find clear evidence for selection in our data, because putatively deleterious variants are purged from our lines more rapidly than presumably neutral variants (Figure 3A), and they are purged more rapidly from high vs. low recombination regions in S6 (Figure 3B). Under this scenario, recombination separates deleterious variants from linked variation, permitting the independent loss of the deleterious variant and allowing neutral diversity to remain (Bersabé et al., 2015). Similarly, subtle relationships were recently discovered within hybrid genomes(Schumer et al., 2018). In the hybrids, high recombination regions retained heterozygosity because recombination broke up incompatibilities that otherwise contribute to hybrid load.

### Outstanding Questions

At least three questions remain. First, what is the mechanism of TE (and knob) removal? One potential explanation is ectopic and/or unequal recombination, which removes TE insertions (Kalendar et al., 2000, Ma and Bennetzen, 2006, Vitte and Bennetzen, 2006). These recombination events can leave a signature of increased numbers of solo to intact LTR elements (Devos et al., 2002), but we found no evidence that TE loss was driven by numerous unequal recombination events among LTRs. It is possible, of course, that unequal recombination caused a small number of large deletion events, with only minor effects on the ratio of solo:intact elements. We nonetheless favor a non-exclusive mechanism for GS loss in this experiment, which is that selection tends to act against the larger haplotype when there is a size difference in a heterozygote. Under this scenario, selfed plants with the best collection of small(er) haplotypes are favored by the selfing process, leading to GS reductions. If true, we expect the resolution of selfing to be a contest between haplotypes, with recombination occasionally reducing interference and combining linked structural variants from different haplotypes onto a single chromosome. Under this model we can make two predictions: i) parental plants of higher heterozygosity and larger differences in size between haplotypes are more likely to lose GS and ii) regions of higher recombination will tend to lose more Mb, due to more efficient selection against large(r) haplotypes. These predictions remain to be tested, underscoring just how little we know about the process of selfing and its effects on genomic variants.

Second, what is the proximal cause of GS loss? The primary effect of selfing is to uncover deleterious recessive mutations, leading to selection against homozygote recessives. But is there a phenotype that drives this selection? GS is known to correlate with several traits, including reproductive rates, growth rates, flowering time, cell sizes and other factors (Rayburn et al., 1994, Knight et al., 2005, Beaulieu et al., 2008, Hu et al., 2011, Tenaillon et al., 2016, Bilinski et al., 2018). Selection on one or several of these diverse characteristics may have occurred during the formation of the inbred lines. However, we cannot find any pattern among our lines that suggest selection was more pronounced on the GSΔ vs. GS_con_ groups. For example, each of the members of GSΔ group (MR01, MR08 and MR18) originated from landraces in the tropical lowlands, and they were bred in lowland tropical nurseries, but the same is true of MR05, MR09, MR11, M22 and MR23, none of which exhibited obvious GS loss.

Finally, what bearing do these results have on broader questions about plant evolution? First, it informs on processes of genome evolution and shows that selection on uncovered, deleterious, putatively recessive alleles can have several effects even over the very-short term. These include removing linked variation in regions of low recombination, purging deleterious alleles in high recombination regions more efficaciously, and hinting that interference between deleterious variants contributes to the retention of heterozygosity, because regions of high heterozygosity tended to be enriched for deleterious variants in S6. Second, this work relates to the finding that indirect selection for recombination modifiers can be favored under selfing (Charlesworth et al., 1979, Roze and Lenormand, 2005). Our results clearly demonstrate that high recombination rates are advantageous for purging genetic load. This relationship may drive the observed trend toward higher chiasmata frequencies in selfing plants compared to outcrossers (Charlesworth and Charlesworth, 1979, Roze and Lenormand, 2005).

## MATERIALS and METHODS

### Plant materials and phenotypic analyses

Our experiment is based on 11 maize landraces (**Table S1**) that were inbred by J. Doebley (U. Wisconsin) and maintained through single-seed descent for several generations (Wills et al., 2013). The parents represent outcrossed landraces of unknown genotype. For each line and generation, one seed was grown and selfed, and the remaining sibling seeds were stored. We grew the sibling seeds in the UC Irvine greenhouses after germination on petri dishes. Ten seeds per cultivar were sown in individual pots on 22 July 2014 and grown in a growth chamber under controlled conditions of 12 h light at 26ºC, 12 h dark at 20ºC, a relative humidity of 70% and 500-600 cal/cm2 of radiation per day. The third and fourth leaves of each plant were harvested when 12-13 cm long and then frozen in liquid nitrogen and stored at −80 ºC. The 11 cultivars, with a subset of 6 plants per cultivar per generation, were grown in four completely randomized blocks, with B73 as the control across blocks. Measures for height were taken on 9, 17, 30 and 45 days after sowing; mortality was also noted throughout the duration. Mortality and growth rates were compared among lines. We estimated the exponential growth rate for each individual and used a one-way ANOVA to test whether the estimate growth rates differed between lines. A logistic regression model was applied to mortality, and a likelihood ratio test was used to compare mortality between lines. We did not measure fitness via fecundity, because none of the lines produced seed under our experimental conditions.

### Flow cytometric data and analyses

To estimate GS, leaf samples were sent to Plant Cytometry Services (Schijndel. Netherlands). Following a previous reference (Diez et al., 2013), flow cytometry used 4’6-diamindino-2-phenylindole (DAPI) staining. Both *Ilex crenata* ‘Fastigiata’ (2C = 2.2pg) and maize B73 (2C = 5.64 pg) (Diez et al., 2013) were employed as internal standards. Three technical replicates were performed for each plant (**Table S2**). To assess whether GS had changed as a consequence of selfing, we performed linear regressions, exponential decay analyses, and Wilcoxon rank sum tests in *R*, combining biological and technical replicates for each time course. Flow cytometeric data were converted to picograms assuming that the maize B73 reference had a value of 5.64 pg/2C (Diez et al., 2013); picograms were translated to Mb assuming 1pg = 978 Mb (Dolezel et al., 2003). To infer a significant trend toward genome loss, we estimated that the probability of loss was 3 lines out of 11 trials (*p*=0.273) and calculated the probability of observing zero GS increases over 11 trials with a two-sided binomial.

### Whole-genome Sequencing and Genomic Composition

We selected six landraces and 33 individuals for whole-genome sequencing (**Table S1**), focusing on the S1 and S6 generations. DNA was extracted from frozen leaf tissue using the QIAGEN DNeasy Plant Mini kit. DNA was multiplexed into libraries with Illumina TruSeq PCR Free kit. The libraries were sequenced on the HiSeq2500 (100 bp read length, paired-end, 2 lanes) in the UCI High Throughput Genomics Facility in 2015 (landraces MR01, MR08, MR18, and MR19) and on the HiSeq3000 (150 bp read length, paired-end, 1 lane) in the UC Davis DNA Technologies Core in 2016 (landraces MR09 and MR22). Individuals were sequenced to an average coverage of ~2.5x per individual (**Table S16**). Note, however, that we had >6x coverage for each generation for each of the lines investigated given the inclusion of siblings.

Sequencing reads were processed by Trimmomatic (v0.35) to remove barcodes and low quality reads (<20), with a minimum read length of 36. Processed reads were mapped simultaneously onto maize genome AGP version 4.37 (AGPv4) (Jiao et al., 2017) and B-specific chromosomal repeats using BWA-MEM (v0.7.12) (Li, 2014). To prevent double counts of a feature, only one of the paired reads was mapped and only the primary alignment was kept for each multi-mapping read, based on Samtools v1.3 (Li et al., 2009).

We counted mapped reads for five annotated genomic components: genes, B-chromosome specific repeats, chromosomal knobs, rDNA and TEs. The annotation features for protein coding genes and for TEs were obtained from the Gramene database on 1/5/17 for B73 AGPv4 (**Table S17**). To annotate regions containing knob (plus CentC) regions and rDNA (plus tDNA) sequences, a series of fasta files (**Table S17**) representing both features were mapped to the v4 genome using blat (v36). The regions of B73 that mapped to either knobs or rDNA were then added to gff files (blattogff v3) for read count analyses. To count reads, all features were merged (bedtools merge v.2.25.0) to avoid double counting (Quinlan, 2014). Bedtools coverage was used to count reads that overlapped at least 90% with each feature. An identical approach was used for W22 annotations (**Table S17**).

We used BUSCO genes to normalize between libraries, on the expectation that these highly conserved genes represent an invariant component of the genome. To identify a conserved set of BUSCO genes, we ran BUSCO (v3) (Simão et al., 2015) on AGPv4. From the resulting set of 1309 BUSCO genes, we eliminated any that appeared to be multi-copy or that overlapped with TE annotations in B73 AGPv4, leaving a final set of 761 genes. A similar procedure in W22 yielded 918 BUSCO genes. In both references, any gene, knob, or rDNA annotation that overlapped with a TE was not considered further. Within any sequencing run, normalized counts for a genomic feature were calculated as the observed number of sequence counts to that feature divided by the total number of counts that mapped to BUSCO genes. To verify that our use of BUSCO genes was accurate, we simulated datasets with BUSCO normalizations based on Chromosome 10 (see below).

Further analyses considered different families and types of TEs. These analyses were performed only in B73. For these, we first identified TEs from the AGPv4 gff file and employed their TE family designations for additional analyses. To examine the ratio of solo LTRs to complete LTRs, we *de novo* annotated Sireviruses based on the MASiVE algorithm (Darzentas et al., 2010). The application of MASiVE produced 22,530 full-length elements with defined boundaries between LTRs and internal regions.

To assess relationships between GS and genomic components, we used both linear regression and ANOVA, using the *lm* and *aov* modules in R (v.3.34). ANOVA p-values were FDR corrected. To estimate the Mb of the genome explained by various component, we: *i*) translated the GS of each plant from pg/2C to Mb, using the conversion rate of 1pg = 978 Mb (Dolezel et al., 2003), *ii*) equated Mb for each individual to the total number of reads mapped to the five genomic components, and *iii*) calculated the number of Mb’s explained per sequencing read. Finally, note that in addition to mapping to our W22 and B73 databases, for completeness we also mapped to a database consisting only of knob repeats, which avoided the complication of reference TE annotations. These analyses also detected a moderate *group*generation* effect (p=0.015) (**Table S18**), suggesting again that knob repeats contribute to GS shifts.

### Testing BUSCO normalization via simulation

To compare counts among individuals, it is important to assess the accuracy of our normalization approach. We tested BUSCO normalization via simulations of TE loss and gain. For the simulations, we used the smallest chromosome 10 for computational efficiency. We randomly removed either 10% or 20% of TEs from the chromosome, duplicated 10% of TEs, or did not change the chromosome. Each treatment was repeated five times with different random TEs removed or gained. The short-read simulator wgsim was used to simulate datasets with ~2x and 10X coverage, mimicking the potential for different coverages among our libraries. For each simulation, reads were mapped to chromosome 10, counted across annotation features (non-BUSCO genes, TEs, knobs and rDNA) and then normalized by dividing by the total counts for BUSCO genes on chromosome 10. We simulated each set of parameter 1000 times. Based on these simulations, we were able to recover the expected decrease in genomic components (Figure S1), but it did not recapitulate genome gain in TEs as accurately. It is likely that the inability to estimate TE gains is a feature of our simulations, because we duplicated TEs as exact, tandem copies of chromosomal TEs, which would lead to systematic undercounting of the duplicated TEs. Nonetheless, our simulations indicate that our normalization approach is sufficient to compare TE loss among datasets with different coverages and different degrees of TE loss.

### Identification of SNPs and deleterious variants

To identify SNPs, paired-end sequencing reads were evaluated for quality using FastQC V0.11.2, and were further processed to remove adapter contamination and low quality bases using Trimmomatic V0.35 (Bolger et al., 2014), with the parameters of LEADING:3, TRAILING:3, SLIDINGWINDOW:4:20, and MINLEN:50. Trimmed reads were then mapped to the B73 reference genome (AGPv4.37; (Jiao et al., 2017) ftp://ftp.ensemblgenomes.org/pub/plants/release-37/fasta/zea_mays) using the MEM algorithm implemented in Burrows-Wheeler Aligner (BWA) V0.7.12 (Li, 2014) with the parameters “-M -k 9 -T 25”. Mapping alignments from one individual were merged using Picard tools V1.96 (http://broadinstitute.github.io/picard/) MergeSamFiles, and potential PCR duplicates were filtered from alignments using SAMtools V1.1 (Li et al., 2009) rmdup. To minimize the number of mismatched bases, local realignment of reads around indels were performed using the Genome Analysis Toolkit (GATK) V3.7 (DePristo et al., 2011) RealignerTargetCreator and IndelRealigner. Only uniquely mapped reads were kept for downstream SNP calling.

To detect SNPs, we used HaplotypeCaller, CombineGVCFs and GenotypeGVCFs from GATK V3.7 (DePristo et al., 2011) separately on each of the six resequenced lines. Variant sites having a minimum phred-scaled confidence threshold 30 and a minimum base quality 20 were considered as SNP candidates. For the SNP set in all samples: *i*) only bi-allelic SNPs were retained, *ii*) genotypes with genotype quality (GQ) score < 5 were assigned as missing, and *iii*) the filtration “QUAL < 30.0, QD < 2.0, MQ < 10.0, DP < 3.0, ReadPosRankSum < −8.0, FS > 30.0” were set to further reduce false positives. A python program parseVCF.py (https://github.com/simonhmartin/genomics_general) was adopted to extract the genotypes of every sample at each SNP site.

We identified putative deleterious SNPs (dSNPs) using SIFT (Kumar et al., 2009), which annotated SNPs as non-coding, synonymous and non-synonymous, based on the gene annotation information in Ensembl (https://plants.ensembl.org). The SIFT database of maize (AGPv3.22) was downloaded from SIFT 4G (http://sift.bii.a-star.edu.sg/sift4g/public/Zea_mays/). Our SNP coordinates were converted to AGPv3 using CrossMap V0.2.7 (Zhao et al., 2014), and then SIFT 4G (Vaser et al., 2016) was launched to compute scores for all converted SNPs. Non-synonymous SNPs (nSNPs) were then predicted as deleterious or tolerated according to their computed SIFT scores. nSNPs having SIFT score < 0.05 were predicted as deleterious; they were considered to be tolerated if they had a normalized probability value ≥ 0.05. For SNPs annotated by SIFT, the derived SNP was inferred using the *Sorghum* genome, based on mapping the raw data from six sorghum varieties from the NCBI short read archive (accession numbers DRR045087, DRR045074, DRR045075, DRR045082, DRR045083 and DRR045081) to the B73 reference. For our analyses, the derived allele was assumed to be the deleterious variant.

### Recombination Data

Crossover data for maize US population were retrieved from (Rodgers-Melnick et al., 2015). The start and end positions of crossover intervals were translated from *Z. mays* B73 AGPv2 to the AGPv4 reference, using CrossMap 0.2.7(Zhao et al., 2014). The number of crossover events in each non-overlapping, 5Mb window was computed as in (Rodgers-Melnick et al., 2015): if a given crossover interval fell over > 1 window, the proportion of the interval present in each window was added to the window crossover counts. Genomic windows were then classified into highly and lowly recombining using the cross-over counts quartiles.

### SNP analyses

We focused only on those SNPs for which the parent could be inferred to be heterozygous – i.e., *H* = 1 in the parent. Operationally, this implied that at least one heterozygote was detected in S1 or that there were two homozygotes with alternative alleles. The derived allele was inferred by comparing SNPs to the Sorghum genome and making the hypothesis that the Sorghum allele is ancestral. SNPs were annotated using SIFT and classified into four categories (see main text). The proportion of the derived allele was computed for each SNP type in each chromosome separately for every line.

A generalized linear model with mixed effects was applied to the proportion of derived allele in each chromosome of every line using the R function *glmer* in the *lme4* package, using the binomial family of tests. Two fixed effects with interaction were considered in the model: the type of SNP as defined by SIFT and the inbreeding generation, see equation (1) below. The line was considered a random effect.

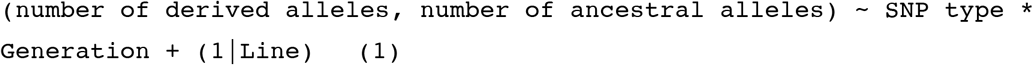

Both fixed effects and their interaction were significant (all p-values < 2.2.10^−16^) using comparison of the fit of model (1) to simpler nested models (removing one effect at a time in model (1)). In order to statistically test whether there was a significant difference between different types of SNPs and/or generations, we computed contrasts with the R package *multcomp*, which automatically corrects for multiple tests.

In order to study the effect of recombination on the proportion of the derived allele, the number of derived and ancestral alleles were summed for each chromosome of every line when considering only highly or lowly recombining genomic windows as previously defined. A similar linear model was then applied, with an additional fixed effect for recombination which interacts with the other two previous fixed effects:

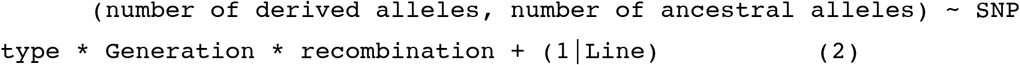

~~~
As previously, all three fixed effects and their interactions were significant when comparing model (2) to simpler nested models (all p-values < 0.007).
~~~

### Heterozygosity Analyses

For each individual, we used sliding windows of 100 SNPs to infer heterozygosity for genomic regions, focusing only on SNPs within genes to avoid potential misalignments due to repetitive elements. Using the set of SNPs inferred to be heterozygous in the parents, the proportion of the major allele *P* was calculated as follows: if a position was homozygous, then the proportion of the major allele was 1. If a position was heterozygous, then the proportion of the major allele was 0.5. The proportion *P* was then averaged across the 100 SNPs of each window for each individual separately to calculate 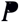. We assumed that the limited number of recombination events in each line over the time course of the experiment did not fully homogenize chromosomes, so that most genomic regions were either heterozygous or homozygous. Based on this approach, the genomic regions that are heterozygous should exhibit a 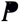 close to 0.5 while genomic regions that are homozygous should have 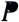 close to 1. Note, however, that real heterozygous loci can be misgenotyped as homozygous to make the 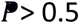. Also, the maize genome contains a high number of duplicated genes, and erroneous mapping of reads from duplicated genes can cause false heterozygous SNPs in homozygous regions (Brandenburg et al., 2017), making 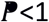 in homozygous regions. Nonetheless, when coverage is high enough to genotype heterozygotes correctly, two peaks of 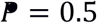 and 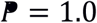 should be observed.

The distribution of 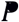 for each line across all individuals and generations is presented in **Figure S11**. Only MR09 and MR22 exhibited the expected two peaks. These two lines have the highest coverage among the set of lines (**Table S16**), and they were therefore the only lines we studied hereafter. Given the distribution of 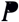 across genomic regions, the *R* package Mclust was used to classify each window of each individual as homozygous or heterozygous (Scrucca et al., 2016) by forcing the number of components to be 2 (G=2). Windows that fell between the two peaks of the 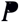 distribution were classified as “uncertain” if the Mclust classification uncertainty was > 0.1 (**Figures S12 and S13**).

For each individual, the heterozygosity status of a region was inferred from the clustering of overlapping sliding windows. The start and end of a heterozygous region were defined by 1) the start of the first window that had the given heterozygosity state and 2) the start of the closest next “uncertain” window. All SNPs inside the region were afterwards considered to be of the inferred heterozygosity type, regardless of genotyping errors. A similar procedure was applied to homozygous regions. Although in principle the categorical status of uncertain regions could be inferred by parsimony arguments, we adopted the conservative approach to discard these blocks of uncertainty from heterozygosity calculations. Heterozygosity levels could then be averaged across individuals of the same line and generation in sliding windows containing 100 SNPs as follows:

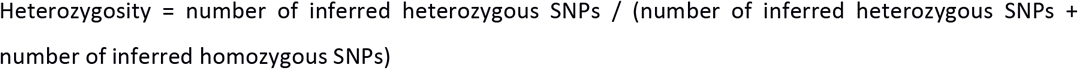

Average heterozygosity levels across individuals were plotted along chromosomes for sliding windows of 100 SNPs that fall within genes (Figure 4). For statistical tests, chromosomes were considered as biologically independent units, owing to the small number of individuals (n=2 or 3). The non-parametric Wilcoxon signed rank test was used to compare the expected heterozygosity with the observed heterozygosity of the ten chromosomes averaged across individuals for each line and generation separately. As a conservative control, this analysis was repeated when considering windows with uncertain heterozygosity in the clustering method as homozygous, instead of discarding them. A similar approach with non overlapping windows of 100 SNPs falling within genes was used to correlate heterozygosity with cross-over number using R lm function. The same non-overlapping windows were used to study the effect of the proportion of nonsynonymous SNPs on heterozygosity using a chi-squared contingency table test with R function chisq.test.

## ACKNOWLEDGEMENTS

We thank four anonymous reviewers for their comments. AM is supported by an EMBO Postdoctoral Fellowship ALTF 775-2017 and by HFSPO fellowship LT000496/2018-L. DS is supported by an NSF Plant Genome Project Fellowship. AB is supported by The Royal Society (Award Numbers UF160222 and RGF\R1\180006). MCS is supported by an NSF Graduate Research Fellowship to UC Davis (1148897). DKS. is supported by a Postdoctoral Fellowship from the National Science Foundation (NSF) Plant Genome Research Program (1609024). JFD is supported by NSF grant IOS 1238014. QL is supported by a National Natural Science Foundation of China grant (no. 31471431) and the Training Program for Outstanding Young Talents of Zhejiang A&F University to QGL. BSG is supported by NSF grants 1542703 and 1655808.

## AUTHOR CONTRIBUTIONS

KR, AM and BSG contributed analyses, ideas and writing. GRJG and QL performed analyses. CMD helped design the experiment, grew plants and measured phenotypes; AB, GRJG, QL, DS, JFD and MS provided materials, data and/or critical ideas. BSG conceived of the project.

## Data and code availability

Sequence data that support the findings of this study have been deposited in NCBI Short Read Archive under project code SRP158803. Custom code used in the analyses is available upon request.

## CITATIONS

Arunkumar R, Ness RW, Wright SI, Barrett SC. 2015. The evolution of selfing is accompanied by reduced efficacy of selection and purging of deleterious mutations. Genetics. 199:817–829.

Barrière A, Yang SP, Pekarek E, Thomas CG, Haag ES, Ruvinsky I. 2009. Detecting heterozygosity in shotgun genome assemblies: Lessons from obligately outcrossing nematodes. Genome Res. 19:470–480.

Beaulieu JM, Leitch IJ, Patel S, Pendharkar A, Knight CA. 2008. Genome size is a strong predictor of cell size and stomatal density in angiosperms. New Phytol. 179:975–986.

Bersabé D, Caballero A, Pérez-Figueroa A, García-Dorado A. 2015. On the Consequences of Purging and Linkage on Fitness and Genetic Diversity. G3 (Bethesda). 6:171–181.

Bilinski P, Albert PS, Berg JJ, Birchler JA, Grote MN, Lorant A, Quezada J, Swarts K, Yang J, Ross-Ibarra J. 2018. Parallel altitudinal clines reveal trends in adaptive evolution of genome size in Zea mays. PLoS Genet. 14:e1007162.

Bolger AM, Lohse M, Usadel B. 2014. Trimmomatic: a flexible trimmer for Illumina sequence data. Bioinformatics. 30:2114–2120.

Bousios A, Kourmpetis YA, Pavlidis P, Minga E, Tsaftaris A, Darzentas N. 2012. The turbulent life of Sirevirus retrotransposons and the evolution of the maize genome: more than ten thousand elements tell the story. Plant J. 69:475–488.

Brandenburg JT, Mary-Huard T, Rigaill G, Hearne SJ, Corti H, Joets J, Vitte C, Charcosset A, Nicolas SD, Tenaillon MI. 2017. Independent introductions and admixtures have contributed to adaptation of European maize and its American counterparts. PLoS Genet. 13:e1006666.

Byers DL, Waller DM. 1999. Do plant populations purge their genetic load? Effects of population size and mating history on inbreeding depression. Annual Review of Ecology and Systematics. 30:479–513.

Charlesworth B, Morgan MT, Charlesworth D. 1993. The effect of deleterious mutations on neutral molecular variation. Genetics. 134:1289–303.

Charlesworth B, Charlesworth, D., Morgan, M.T. 1990. Genetic loads and estimates of mutation rates in highly inbred plant populations. Nature. 347:380–382.

Charlesworth D, Charlesworth B, Strobeck C. 1979. Selection for recombination in partially self-fertilizing populations. Genetics. 93:237–244.

Charlesworth D, Willis JH. 2009. The genetics of inbreeding depression. Nat Rev Genet. 10:783–796.

Charlesworth D, Charlesworth B. 1979. The eovlutionary genetics of sexual systems in flowering plants. Proc R Soc Lond B Biol Sci. 205:513–530.

Charlesworth D, Wright SI. 2001. Breeding systems and genome evolution. Curr Opin Genet Dev. 11:685–690.

Chia JM, Song C, Bradbury PJ et al. 2012. Maize HapMap2 identifies extant variation from a genome in flux. Nat Genet. 44:803–807.

Crnokrak P, Barrett SC. 2002. Perspective: purging the genetic load: a review of the experimental evidence. Evolution. 56:2347–2358.

Crow JF. 2008. Mid-century controversies in population genetics. Annu Rev Genet. 42:1–16.

Cullis CA. 2005. Mechanisms and control of rapid genomic changes in flax. Ann Bot. 95:201–206.

Darwin C. 1876. The effects of self and cross fertilization in the vegetable kingdom. London: John Murray.

Darzentas N, Bousios A, Apostolidou V, Tsaftaris AS. 2010. MASiVE: Mapping and Analysis of Sirevirus Elements in plant genome sequences. Bioinformatics. 26:2452–2454.

DePristo MA, Banks E, Poplin R et al. 2011. A framework for variation discovery and genotyping using next-generation DNA sequencing data. Nat Genet. 43:491–498.

Devos KM, Brown JK, Bennetzen JL. 2002. Genome size reduction through illegitimate recombination counteracts genome expansion in Arabidopsis. Genome Res. 12:1075–9.

Diez CM, Gaut BS, Meca E, Scheinvar E, Montes-Hernandez S, Eguiarte LE, Tenaillon MI. 2013. Genome size variation in wild and cultivated maize along altitudinal gradients. New Phytol. doi: 10.1111/nph.12247

Diez CM, Meca E, Tenaillon MI, Gaut BS. 2014. Three Groups of Transposable Elements with Contrasting Copy Number Dynamics and Host Responses in the Maize (Zea mays ssp. mays) Genome. PLoS Genet. 10:e1004298.

Dolezel J, Bartos J, Voglmayr H, Greilhuber J. 2003. Nuclear DNA content and genome size of trout and human. Cytometry A. 51:127–8; author reply 129.

Fierst JL, Willis JH, Thomas CG, Wang W, Reynolds RM, Ahearne TE, Cutter AD, Phillips PC. 2015. Reproductive Mode and the Evolution of Genome Size and Structure in Caenorhabditis Nematodes. PLoS Genet. 11:e1005323.

Fisher RA. 1941. Average excess and average effect of a gene substitution. Annals of Human Genetics. 11:53–63.

Gore MA, Chia JM, Elshire RJ et al. 2009. A first-generation haplotype map of maize. Science. 326:1115–1117.

Govindaraju DIDDAHALLYR, Cullis CHRISTOPHERA. 1991. Modulation of genome size in plants: the influence of breeding systems and neighbourhood size. Evolutionary Trends in Plants (United Kingdom).

Hedrick PW. 1994. Purging inbreeding depression and the probability of extinction: full-sib mating. Heredity (Edinb). 73:363–372.

Hedrick PW, Garcia-Dorado A. 2016. Understanding Inbreeding Depression, Purging, and Genetic Rescue. Trends Ecol Evol. 31:940–952.

Hedrick PW, Hellsten U, Grattapaglia D. 2016. Examining the cause of high inbreeding depression: analysis of whole-genome sequence data in 28 selfed progeny of Eucalyptus grandis. New Phytol. 209:600–611.

Hill WG, Robertson A. 1966. The effect of linkage on limits to artificial selection. Genet Res. 8:269–94.

Hollister JD, Gaut BS. 2009. Epigenetic silencing of transposable elements: A trade-off between reduced transposition and deleterious effects on neighboring gene expression. Genome Res. 19:1419–1428.

Hollister JD, Smith LM, Guo YL, Ott F, Weigel D, Gaut BS. 2011. Transposable elements and small RNAs contribute to gene expression divergence between Arabidopsis thaliana and Arabidopsis lyrata. Proc Natl Acad Sci U S A.

Hu TT, Pattyn P, Bakker EG et al. 2011. The Arabidopsis lyrata genome sequence and the basis of rapid genome size change. Nat Genet. 43:476–481.

Jian Y, Xu C, Guo Z, Wang S, Xu Y, Zou C. 2017. Maize (Zea mays L.) genome size indicated by 180-bp knob abundance is associated with flowering time. Sci Rep. 7:5954.

Jiao Y, Peluso P, Shi J et al. 2017. Improved maize reference genome with single-molecule technologies. Nature. 546:524–527.

Kalendar R, Tanskanen J, Immonen S, Nevo E, Schulman AH. 2000. Genome evolution of wild barley (Hordeum spontaneum) by BARE-1 retrotransposon dynamics in response to sharp microclimatic divergence. Proc Natl Acad Sci U S A. 97:6603–6607.

Kardos M, Taylor HR, Ellegren H, Luikart G, Allendorf FW. 2016. Genomics advances the study of inbreeding depression in the wild. Evol Appl. 9:1205–1218.

Knight CA, Molinari NA, Petrov DA. 2005. The large genome constraint hypothesis: Evolution, ecology and phenotype. Calif Polytech State Univ San Luis Obispo, Dept Biol Sci, San Luis Obispo, CA 93407 USA:

Kumar P, Henikoff S, Ng PC. 2009. Predicting the effects of coding non-synonymous variants on protein function using the SIFT algorithm. Nat Protoc. 4:1073–1081.

Lande R, Schemske DW. 1985. The evolution of self-fertilization and inbreeding depression in plants. I. Genetic models. Evolution. 39:24–40.

Lee YCG, Karpen GH. 2017. Pervasive epigenetic effects of Drosophila euchromatic transposable elements impact their evolution. Elife. 6

Li H. 2014. Toward better understanding of artifacts in variant calling from high-coverage samples. Bioinformatics. 30:2843–2851.

Li H, Handsaker B, Wysoker A, Fennell T, Ruan J, Homer N, Marth G, Abecasis G, Durbin R. 2009. The Sequence Alignment/Map format and SAMtools. Bioinformatics. 25:2078–2079.

Liu Q, Zhou Y, Morrell PL, Gaut BS. 2017. Deleterious Variants in Asian Rice and the Potential Cost of Domestication. Mol Biol Evol. 34:908–924.

Long Q, Rabanal FA, Meng D et al. 2013. Massive genomic variation and strong selection in Arabidopsis thaliana lines from Sweden. Nat Genet. 45:884–890.

Lyu H, He Z, Wu CI, Shi S. 2018. Convergent adaptive evolution in marginal environments: unloading transposable elements as a common strategy among mangrove genomes. New Phytol. 217:428–438.

Ma J, Bennetzen JL. 2006. Recombination, rearrangement, reshuffling, and divergence in a centromeric region of rice. Proc Natl Acad Sci U S A. 103:383–388.

McMullen MD, Kresovich S, Villeda HS et al. 2009. Genetic properties of the maize nested association mapping population. Science. 325:737–740.

Mezmouk S, Ross-Ibarra J. 2014. The pattern and distribution of deleterious mutations in maize. G3 (Bethesda). 4:163–171.

Morran LT, Ohdera AH, Phillips PC. 2010. Purging deleterious mutations under self fertilization: paradoxical recovery in fitness with increasing mutation rate in Caenorhabditis elegans. PLoS One. 5:e14473.

Morran LT, Parmenter MD, Phillips PC. 2009. Mutation load and rapid adaptation favour outcrossing over self-fertilization. Nature. 462:350–352.

Mroczek RJ, Melo JR, Luce AC, Hiatt EN, Dawe RK. 2006. The maize Ab10 meiotic drive system maps to supernumerary sequences in a large complex haplotype. Genetics. 174:145–154.

Ng PC, Henikoff S. 2003. SIFT: Predicting amino acid changes that affect protein function. Nucleic Acids Res. 31:3812–3814.

Ohta T. 1971. Associative overdominance caused by linked detrimental mutations. Genet Res. 18:277–286.

Price HJ. 1976. Evolution of DNA content in higher plants. The Botanical Review. 42:27.

Quadrana L, Bortolini Silveira A, Mayhew GF, LeBlanc C, Martienssen RA, Jeddeloh JA, Colot V. 2016. The Arabidopsis thaliana mobilome and its impact at the species level. Elife. 5

Quinlan AR. 2014. BEDTools: The Swiss-Army Tool for Genome Feature Analysis. Curr Protoc Bioinformatics. 47:11.12.1–34.

Randolph LF. 1941. Genetic characteristics of the B chromosomes in maize. Genetics. 26:608–631.

Rayburn AL, Dudley JW, Biradar DP. 1994. Selection for early flowering results in simultaneous selection for reduced nuclear-DNA content in maize. Plant Breeding. 112:318–322.

Rodgers-Melnick E, Bradbury PJ, Elshire RJ, Glaubitz JC, Acharya CB, Mitchell SE, Li C, Li Y, Buckler ES. 2015. Recombination in diverse maize is stable, predictable, and associated with genetic load. Proc Natl Acad Sci U S A. 112:3823–3828.

Ross-Ibarra J, Tenaillon M, Gaut BS. 2009. Historical divergence and gene flow in the genus zea. Genetics. 181:1399–1413.

Roze D, Lenormand T. 2005. Self-fertilization and the evolution of recombination. Genetics. 170:841–857.

Schnable PS, Springer NM. 2013. Progress toward understanding heterosis in crop plants. Annu Rev Plant Biol. 64:71–88.

Schnable PS, Ware D, Fulton RS et al. 2009. The B73 maize genome: complexity, diversity, and dynamics. Science. 326:1112–1115.

Schultz ST, Willis JH. 1995. Individual variation in inbreeding depression: the roles of inbreeding history and mutation. Genetics. 141:1209–1223.

Schumer M, Xu C, Powell DL et al. 2018. Natural selection interacts with recombination to shape the evolution of hybrid genomes. Science. 360:656–660.

Scrucca L, Fop M, Murphy TB, Raftery AE. 2016. mclust 5: Clustering, Classification and Density Estimation Using Gaussian Finite Mixture Models. R J. 8:289–317.

Simão FA, Waterhouse RM, Ioannidis P, Kriventseva EV, Zdobnov EM. 2015. BUSCO: assessing genome assembly and annotation completeness with single-copy orthologs. Bioinformatics. 31:3210–3212.

Smarda P, Horova L, Bures P, Hralova I, Markova M. 2010. Stabilizing selection on genome size in a population of Festuca pallens under conditions of intensive intraspecific competition. New Phytol. 187:1195–1204.

Springer NM, Anderson SN, Andorf CM et al. 2018. The maize W22 genome provides a foundation for functional genomics and transposon biology. Nat Genet.

Springer NM, Stupar RM. 2007. Allelic variation and heterosis in maize: how do two halves make more than a whole? Genome Res. 17:264–275.

Sun S, Zhou Y, Chen J et al. 2018. Extensive intraspecific gene order and gene structural variations between Mo17 and other maize genomes. Nat Genet.

Takebayashi N, Morrell PL. 2001. Is self-fertilization an evolutionary dead end? Revisiting an old hypothesis with genetic theories and a macroevolutionary approach. Am J Bot. 88:1143–1150.

Tenaillon MI, Hollister JD, Gaut BS. 2010. A triptych of the evolution of plant transposable elements. Trends Plant Sci. 15:471–478.

Tenaillon MI, Hufford MB, Gaut BS, Ross-Ibarra J. 2011. Genome Size and Transposable Element Content as Determined by High-Throughput Sequencing in Maize and Zea luxurians. Genome Biol Evol. 3:219–229.

Tenaillon MI, Manicacci D, Nicolas SD, Tardieu F, Welcker C. 2016. Testing the link between genome size and growth rate in maize. PeerJ. 4:e2408.

Thornton KR, Foran AJ, Long AD. 2013. Properties and modeling of GWAS when complex disease risk is due to non-complementing, deleterious mutations in genes of large effect. PLoS Genet. 9:e1003258.

Vaser R, Adusumalli S, Leng SN, Sikic M, Ng PC. 2016. SIFT missense predictions for genomes. Nat Protoc. 11:1–9.

Vitte C, Bennetzen JL. 2006. Analysis of retrotransposon structural diversity uncovers properties and propensities in angiosperm genome evolution. Proc Natl Acad Sci U S A. 103:17638–17643.

Weller SG, Sakai AK, Thai DA, Tom J, Rankin AE. 2005. Inbreeding depression and heterosis in populations of Schiedea viscosa, a highly selfing species. J Evol Biol. 18:1434–1444.

Wills DM, Whipple CJ, Takuno S, Kursel LE, Shannon LM, Ross-Ibarra J, Doebley JF. 2013. From many, one: genetic control of prolificacy during maize domestication. PLoS Genet. 9:e1003604.

Wright SI, Kalisz S, Slotte T. 2013. Evolutionary consequences of self-fertilization in plants. Proc Biol Sci. 280:20130133.

Wright SI, Ness RW, Foxe JP, Barrett SCH. 2008. Genomic consequences of outcrossing and selfing in plants. International Journal of Plant Sciences. 169:105–118.

Yamakake K. 1976. Cytological studies in maize (Zea mays L.) and teosinte (Zea mexicana (Schrader) Kuntze) in relation to their origin and evolution. Bull Mass Agric Exp Stat.

Yin D, Schwarz EM, Thomas CG, Felde RL, Korf IF, Cutter AD, Schartner CM, Ralston EJ, Meyer BJ, Haag ES. 2018. Rapid genome shrinkage in a self-fertile nematode reveals sperm competition proteins. Science. 359:55–61.

Zhao H, Sun Z, Wang J, Huang H, Kocher JP, Wang L. 2014. CrossMap: a versatile tool for coordinate conversion between genome assemblies. Bioinformatics. 30:1006–1007.

