## Supplemental Figures and Tables for "The Genomics of Selfing in Maize (*Zea mays* ssp. *mays*): Catching Purging in the Act"

***** co first-authors

**Table S1**: Sampling of individuals for each lines for flow cytometry or for WGS resequencing.

| **Line** | **Landrace** | **Flow Cytometry Gen. (No.)** | **WGS Gen. (No.)** |
|---|---|---|---|
| MR01 | Araguito | 1(3), 2(1),4(3), 5(1), 6(1) | 1(3), 5(1), 6(1) |
| MR05 | Cateto | 1(3), 6(3) | -- |
| MR08 | Costeno | 1(3), 2(3), 3(3), 4(3), 6(3) | 1(3), 6(3) |
| MR09 | Cravo Rioganense | 1(2), 2(3), 6(3) | 1(2), 6(2) |
| MR11 | Cuban Flint | 1(2), 2(3), 5(3), 6(3) | -- |
| MR13 | Hickory King | 1(2), 7(1), 8(1) | -- |
| MR14 | Long Fellow Flint | 1(3), 5(3), 6(1) | -- |
| MR18 | Reventador | 1(3), 2(3), 3(3), 4(3), 5(2), 6(3) | 1(3), 6(3) |
| MR19 | Santo Domingo | 1(3), 7(3) | 1(3), 6(3) |
| MR22 | Tuxpeno | 1(3), 6(3) | 1(3), 6(3) |
| MR23 | Zapalote Chico | 1(1), 2(3),6(3) | -- |
| **TOTAL** | | 96 | 33 |

**Table S2:** Raw data from flow cytometry, representing three technical replicates per plant.

| Plant | Landrace | Generation | Rep1_pg | Rep2_pg | Rep3_pg | Avg |
|---|---|---|---|---|---|---|
| 1 | MR09 | 1 | 5.939 | 5.939 | 5.901 | 5.926 |
| 2 | MR09 | 2 | 6.424 | 6.462 | 6.163 | 6.350 |
| 5 | MR09 | 6 | 6.126 | 5.976 | 5.827 | 5.976 |
| 6 | MR01 | 1 | 7.283 | 7.358 | 7.283 | 7.308 |
| 9 | MR01 | 4 | 6.542 | 6.618 | 6.768 | 6.643 |
| 12 | MR05 | 1 | 6.014 | 5.976 | 6.051 | 6.014 |
| 17 | MR05 | 6 | 5.901 | 5.901 | 6.014 | 5.939 |
| 18 | MR11 | 1 | 6.014 | 5.939 | 6.163 | 6.038 |
| 19 | MR11 | 2 | 6.051 | 6.014 | 6.051 | 6.038 |
| 22 | MR11 | 5 | 5.715 | 5.752 | 5.677 | 5.715 |
| 23 | MR11 | 6 | 6.200 | 6.051 | 6.126 | 6.126 |
| 24 | MR19 | 1 | 7.022 | 7.097 | 6.947 | 7.022 |
| 30 | MR19 | 7 | 7.171 | 7.134 | 7.097 | 7.134 |
| 31 | MR22 | 1 | 6.088 | 6.126 | 5.976 | 6.063 |
| 36 | MR22 | 6 | 6.088 | 6.126 | 6.014 | 6.076 |
| 37 | MR13 | 1 | 5.789 | 5.603 | 5.827 | 5.740 |
| 43 | MR13 | 8 | 5.827 | 5.939 | 5.901 | 5.889 |
| 44 | MR14 | 1 | 5.416 | 5.491 | 5.528 | 5.478 |
| 48 | MR14 | 5 | 5.528 | 5.603 | 5.677 | 5.603 |
| 50 | MR08 | 1 | 7.059 | 6.835 | 6.910 | 6.935 |
| 51 | MR08 | 2 | 6.881 | 6.994 | 6.994 | 6.956 |
| 52 | MR08 | 3 | 6.843 | 7.106 | 6.994 | 6.981 |
| 53 | MR08 | 4 | 6.843 | 6.768 | 6.806 | 6.806 |
| 54 | MR08 | 6 | 6.686 | 6.499 | 6.648 | 6.611 |
| 56 | MR23 | 2 | 6.648 | 6.686 | 6.462 | 6.599 |
| 59 | MR23 | 6 | 6.686 | 6.499 | 6.536 | 6.574 |
| 60 | MR18 | 1 | 6.275 | 6.424 | 6.312 | 6.337 |
| 61 | MR18 | 2 | 6.317 | 6.242 | 6.242 | 6.267 |
| 62 | MR18 | 3 | 6.054 | 6.054 | 6.129 | 6.079 |
| 63 | MR18 | 4 | 6.166 | 6.129 | 6.129 | 6.141 |
| 64 | MR18 | 5 | 6.204 | 6.166 | 5.941 | 6.104 |
| 65 | MR18 | 6 | 6.088 | 6.163 | 6.126 | 6.126 |
| 72 | MR11 | 2 | 6.275 | 5.901 | 6.088 | 6.088 |
| 75 | MR11 | 5 | 6.238 | 6.200 | 6.424 | 6.287 |
| 76 | MR11 | 6 | 6.014 | 6.051 | 5.939 | 6.001 |
| 77 | MR14 | 1 | 5.453 | 5.416 | 5.491 | 5.453 |
| 81 | MR14 | 5 | 5.491 | 5.453 | 5.528 | 5.491 |
| 83 | MR18 | 1 | 6.499 | 6.462 | 6.312 | 6.424 |

| | | | | | | |
|---|---|---|---|---|---|---|
| 84 | MR18 | 2 | 6.091 | 6.467 | 6.242 | 6.267 |
| 85 | MR18 | 3 | 6.091 | 6.166 | 6.091 | 6.116 |
| 86 | MR18 | 4 | 6.091 | 6.091 | 6.129 | 6.104 |
| 88 | MR18 | 6 | 5.827 | 6.014 | 5.939 | 5.926 |
| 94 | MR19 | 1 | 6.686 | 6.985 | 6.947 | 6.873 |
| 100 | MR19 | 7 | 7.171 | 7.209 | 7.134 | 7.171 |
| 101 | MR13 | 1 | 5.827 | 5.901 | 5.901 | 5.877 |
| 108 | MR09 | 1 | 5.901 | 5.752 | 5.789 | 5.814 |
| 109 | MR09 | 2 | 5.491 | 5.640 | 5.677 | 5.603 |
| 112 | MR09 | 6 | 5.901 | 5.752 | 5.789 | 5.814 |
| 113 | MR05 | 1 | 5.491 | 5.640 | 5.677 | 5.603 |
| 118 | MR05 | 6 | 5.827 | 5.752 | 5.789 | 5.789 |
| 119 | MR23 | 1 | 6.387 | 6.387 | 6.387 | 6.387 |
| 120 | MR23 | 2 | 6.387 | 6.387 | 6.387 | 6.387 |
| 123 | MR23 | 6 | 6.462 | 6.499 | 6.350 | 6.437 |
| 124 | MR01 | 1 | 7.433 | 7.395 | 7.395 | 7.408 |
| 125 | MR01 | 2 | 6.768 | 6.806 | 6.881 | 6.818 |
| 127 | MR01 | 4 | 6.918 | 6.881 | 6.730 | 6.843 |
| 129 | MR01 | 6 | 6.611 | 6.723 | 6.798 | 6.711 |
| 130 | MR08 | 1 | 6.835 | 6.873 | 6.798 | 6.835 |
| 131 | MR08 | 2 | 7.069 | 7.144 | 7.031 | 7.081 |
| 132 | MR08 | 3 | 6.730 | 6.768 | 6.881 | 6.793 |
| 133 | MR08 | 4 | 6.693 | 6.655 | 6.618 | 6.655 |
| 134 | MR08 | 6 | 6.723 | 6.761 | 6.723 | 6.736 |
| 135 | MR22 | 1 | 6.088 | 5.976 | 6.014 | 6.026 |
| 140 | MR22 | 6 | 6.126 | 6.088 | 6.200 | 6.138 |
| 141 | MR18 | 1 | 6.947 | 6.947 | 6.947 | 6.947 |
| 142 | MR18 | 2 | 6.054 | 6.091 | 6.091 | 6.079 |
| 143 | MR18 | 3 | 6.091 | 6.129 | 6.129 | 6.116 |
| 144 | MR18 | 4 | 6.054 | 6.016 | 6.091 | 6.054 |
| 145 | MR18 | 5 | 5.978 | 6.016 | 6.054 | 6.016 |
| 146 | MR18 | 6 | 6.088 | 6.126 | 6.088 | 6.101 |
| 152 | MR05 | 1 | 5.901 | 5.901 | 5.939 | 5.914 |
| 157 | MR05 | 6 | 6.014 | 6.051 | 6.014 | 6.026 |
| 158 | MR01 | 1 | 7.022 | 7.022 | 7.134 | 7.059 |
| 161 | MR01 | 4 | 6.761 | 6.761 | 6.761 | 6.761 |
| 162 | MR01 | 5 | 6.723 | 6.761 | 6.648 | 6.711 |
| 164 | MR22 | 1 | 6.200 | 6.088 | 6.163 | 6.150 |
| 169 | MR22 | 6 | 6.126 | 6.051 | 6.126 | 6.101 |
| 175 | MR13 | 7 | 5.715 | 5.827 | 5.827 | 5.789 |
| 177 | MR11 | 1 | 6.200 | 6.163 | 6.014 | 6.126 |
| 178 | MR11 | 2 | 5.939 | 6.051 | 6.014 | 6.001 |
| 181 | MR11 | 5 | 6.163 | 6.051 | 6.126 | 6.113 |

| | | | | | | |
|---|---|---|---|---|---|---|
| 182 | MR11 | 6 | 5.901 | 5.976 | 5.789 | 5.889 |
| 183 | MR08 | 1 | 6.723 | 6.798 | 6.761 | 6.761 |
| 184 | MR08 | 2 | 6.768 | 6.918 | 6.655 | 6.781 |
| 185 | MR08 | 3 | 6.806 | 6.881 | 6.806 | 6.831 |
| 186 | MR08 | 4 | 6.918 | 6.956 | 6.843 | 6.906 |
| 187 | MR08 | 6 | 6.611 | 6.499 | 6.723 | 6.611 |
| 188 | MR14 | 1 | 5.416 | 5.528 | 5.528 | 5.491 |
| 192 | MR14 | 5 | 5.453 | 5.491 | 5.901 | 5.615 |
| 193 | MR14 | 6 | 5.453 | 5.341 | 5.416 | 5.403 |
| 195 | MR23 | 2 | 6.462 | 6.611 | 6.499 | 6.524 |
| 198 | MR23 | 6 | 6.350 | 6.499 | 6.424 | 6.424 |
| 200 | MR09 | 2 | 5.752 | 5.827 | 5.901 | 5.827 |
| 203 | MR09 | 6 | 5.752 | 5.789 | 5.715 | 5.752 |
| 204 | MR19 | 1 | 7.321 | 7.283 | 7.246 | 7.283 |
| 210 | MR19 | 7 | 7.171 | 7.059 | 6.910 | 7.047 |

**Table S3:** Table of results from Wilcoxson rank-sum tests that compare flow cytometric estimates of GS between generation S1 and a later generation. Bolded values represent significance at $\alpha < 0.05$ after Bonferroni correction.

| | Gens | Wilcoxson p-value |
|---|---|---|
| **MRO1** | **1,4** | **0.004** |
| MR05 | 1,6 | 0.532 |
| **MR08** | **1,6** | **0.001** |
| MR09 | 1,6 | 0.512 |
| MR11 | 1,6 | 0.313 |
| MR13 | 1,8 | 0.229 |
| MR14 | 1,5 | 0.105 |
| **MR18** | **1,6** | **0.004** |
| MR19 | 1,7 | 0.546 |
| MR22 | 1,6 | 0.470 |
| MR23 | 1,6 | 0.189 |

**Table S4:** Results for tests of negative GS slopes over time based on flow cytometric estimates, using both a linear model of loss and a model of exponential decay. Bolded values are significant $\alpha < 0.05$ after Bonferroni correction. Among the three significant lines, the exponential decay model fits better for lines MR01 and MR18, based on the Akaike Information Criterion (AIC). 'Estimate' represents the estimate of the slope of the linear model or the exponential parameter.

| | Linear Model | | | Exponential Decay | | |
|---|---|---|---|---|---|---|
| | Estimate | p-value | AIC | Estimate | p-value | AIC |
| **MRO1** | **-0.032590** | **1.03e-06** | **-84.21** | **0.40244** | **1.58e-06** | **-93.81** |
| MR05 | 0.004000 | 0.329 | -59.02 | -0.03842 | 0.334 | -59.02 |
| **MR08** | **-0.012117** | **6.65e-05** | **-179.08** | **0.14906** | **0.000684** | **-177.55** |
| MR09 | -0.002844 | 0.628 | -61.99 | 0.02674 | 0.642 | -61.98 |
| MR11 | -0.002942 | 0.444 | -109.03 | 0.03034 | 0.443 | -109.03 |
| MR13 | 0.001579 | 0.495 | -50.43 | -0.02764 | 0.48340 | -50.46 |
| MR14 | 0.002267 | 0.491 | -81.78 | -0.04645 | 0.5496 | -81.72 |
| **MR18** | **-0.023118** | **1.21e-06** | **-154.42** | **0.38602** | **2.23e-08** | **-167.37** |
| MR19 | 0.002593 | 0.442 | -59.29 | -0.02410 | 0.445 | -59.29 |
| MR22 | 0.001333 | 0.443 | -89.69 | -0.043022 | 0.44925 | -89.69 |
| MR23 | 0.000991 | 0.743 | -85.18 | -0.02662 | 0.764 | -85.16 |

**Table S5:** Estimates of the variance components and p-values with normalized read count data based on an ANOVA model and the B73 reference dataset. Each of the five genomic components (TEs, genes, knob-repeats, B chromosome specific repeats and rDNA) was tested individually for the Entire B73 Genome.

| | Group | Landrace | Generation | Group X Gen | Race X Gen |
|---|---|---|---|---|---|
| TE_v[1] | 14.722 | 70.655 | 2.8234 | 5.415 | 0.651 |
| TE_p[2] | **2.37e-06** | **2.91-10** | **1.80e-02** | **1.32e-03** | 8.36e-01 |
| gene_v | 1.463 | 21.487 | 4.849 | 0.0169 | 18.515 |
| gene_p | 5.97e-01 | 2.50e-01 | 2.89e-01 | 1.00 | 2.89e-01 |
| knob_v | 35.604 | 56.212 | 0.444 | 0.497 | 2.540 |
| knob_p | **2.91e-10** | **2.91e-10** | 0.289 | 0.289 | 0.116 |
| bChr_v | 7.020 | 25.797 | 7.269 | 6.757 | 25.529 |
| bChr_p | 0.085 | **0.021** | 0.085 | 0.085 | **0.0219** |
| rDNA_v | 1.996 | 40.489 | 2.272 | 1.528 | 13.467 |
| rDNA_p | 0.456 | **0.018** | 0.433 | 0.521 | 0.289 |

[1] The suffix v refers to the percentage of the total variance explained.
[2] The suffix p refers to probability. Bolded values are significant at $p$ = 0.05 after FDR correction for all tests in the Table.

**Table S6:** Estimates of the average Mb attributable to each of the five genomic components, based on mapping to the annotated B73 reference genome.

| Landrace | Generation | GS[1] | Genes[2] | TEs[2] | Knobs[2] | rDNA[2] | B-Chrom[2] |
|---|---|---|---|---|---|---|---|
| MR01 | S1 | 7.259 | 339.999 | 6750.690 | 0.508 | 7.645 | 0.013 |
| MR01 | S6 | 6.764 | 338.668 | 6269.190 | 0.507 | 7.237 | 0.012 |
| | Diff (S1-S6) | | 1.331 | 481.500 | 0.001 | 0.407 | 0.001 |
| MR08 | S1 | 6.844 | 327.931 | 6357.124 | 0.509 | 7.376 | 0.033 |
| MR08 | S6 | 6.653 | 339.102 | 6158.515 | 0.508 | 8.070 | 0.073 |
| | Diff (S1-S6) | | -11.171 | 198.609 | 0.001 | -0.693 | -0.040 |
| MR09 | S1 | 5.831 | 345.315 | 5388.369 | 0.378 | 7.117 | 0.005 |
| MR09 | S6 | 5.848 | 334.501 | 5377.505 | 0.380 | 6.471 | 0.004 |
| | Diff (S1-S6) | | 10.814 | 10.864 | -0.001 | 0.646 | 0.001 |
| MR18 | S1 | 6.192 | 340.976 | 6063.259 | 0.497 | 9.702 | 10.660 |
| MR18 | S6 | 6.333 | 311.376 | 5598.424 | 0.454 | 7.434 | 0.055 |
| | Diff (S1-S6) | | 29.600 | 464.835 | 0.043 | 2.268 | 10.605 |
| MR19 | S1 | 6.665 | 331.432 | 6563.089 | 0.514 | 8.990 | 0.007 |
| MR19 | S6 | 7.117 | 310.996 | 6571.233 | 0.483 | 9.138 | 0.006 |
| | Diff (S1-S6) | | 20.436 | -8.144 | 0.031 | -0.148 | 0.002 |
| MR22 | S1 | 6.088 | 338.116 | 5626.868 | 0.405 | 7.137 | 0.009 |
| MR22 | S6 | 6.101 | 339.521 | 5637.490 | 0.392 | 7.297 | 0.013 |
| | Diff (S1-S6) | | -1.404 | -10.621 | 0.012 | -0.159 | -0.004 |

[1] Genome Size (GS) estimates are in pg/2c. For MR19, GS estimates are from three S7 plants but sequence data were from three S6 plants.
[2] Values are estimates of Mb attributable to each genomic component, averaged across individuals within each landrace and generation.

**Table S7:** Estimates of the variance components and p-values with normalized read count data based on an ANOVA model and mapping to the W22 reference. Each of the five genomic components (TEs, genes, knob-repeats, B chromosome specific repeats and rDNA) was tested individually for the Entire W22 Genome.

| | Group | Landrace | Generation | Group X Gen | Race X Gen |
|---|---|---|---|---|---|
| TE_v[1] | 14.705 | 69.346 | 3.460 | 6.137 | 1.115 |
| TE_p[2] | **1.58E-07** | **1.24E-10** | **5.36E-03** | **3.94E-04** | 5.35E-01 |
| gene_v | 6.646 | 61.360 | 3.963 | 4.388 | 7.147 |
| gene_p | 2.29E-02 | **5.84E-06** | 6.67E-02 | 6.41E-02 | 1.59E-01 |
| knob_v | 5.122 | 85.143 | 1.002 | 2.727 | 0.745 |
| knob_p | **9.34E-04** | **3.51E-11** | 1.04E-01 | **1.28E-02** | 7.47E-01 |
| bChr_v | 6.996 | 25.938 | 7.351 | 6.825 | 25.711 |
| bChr_p | **6.41E-02** | **1.69E-02** | 6.41E-02 | 6.41E-02 | **1.69E-02** |
| rDNA_v | 0.0458 | 11.597 | 4.325 | 0.716 | 18.815 |
| rDNA_p | 1.00E+00 | 6.25E-01 | **3.73E-01** | 7.93E-01 | **3.62E-01** |

[1] The suffix v refers to the percentage of the total variance explained.

[2] The suffix p refers to probability. Bolded values are significant at *p* = 0.05 after FDR correction for all tests in the Table.

**Table S8:** Estimates of the average Mb attributable to each of the five genomic components, based on mapping to the annotated W22 reference genome.

| Landrace | Generation | GS[1] | Genes[2] | TEs[2] | Knobs[2] | rDNA[2] | B-Chrom[2] |
|---|---|---|---|---|---|---|---|
| MR01 | S1 | 7.259 | 298.260 | 6205.137 | 584.873 | 10.575 | 0.011 |
| MR01 | S6 | 6.764 | 293.441 | 5863.664 | 448.462 | 10.036 | 0.010 |
| | Diff (S1-S6) | | 4.819 | 341.473 | 136.411 | 0.538 | 0.001 |
| MR08 | S1 | 6.844 | 291.407 | 5924.244 | 467.867 | 9.434 | 0.023 |
| MR08 | S6 | 6.653 | 293.002 | 5794.143 | 408.503 | 10.571 | 0.049 |
| | Diff (S1-S6) | | -1.595 | 130.101 | 59.364 | -1.138 | -0.026 |
| MR09 | S1 | 5.831 | 302.420 | 5285.736 | 141.403 | 11.622 | 0.003 |
| MR09 | S6 | 5.848 | 300.623 | 5267.027 | 140.122 | 11.086 | 0.003 |
| | Diff (S1-S6) | | 1.798 | 18.709 | 1.281 | 0.536 | 0.000 |
| MR18 | S1 | 6.192 | 299.143 | 5799.792 | 302.795 | 12.647 | 10.716 |
| MR18 | S6 | 6.333 | 296.334 | 5385.928 | 225.797 | 9.646 | 0.037 |
| | Diff (S1-S6) | | 2.809 | 413.864 | 76.998 | 3.001 | 10.679 |
| MR19 | S1 | 6.665 | 302.115 | 6041.574 | 549.868 | 10.467 | 0.007 |
| MR19 | S6 | 7.117 | 285.282 | 5952.045 | 644.749 | 9.777 | 0.004 |
| | Diff (S1-S6) | | 16.834 | 89.529 | -94.881 | 0.691 | 0.004 |
| MR22 | S1 | 6.088 | 301.452 | 5446.861 | 213.602 | 10.613 | 0.008 |
| MR22 | S6 | 6.101 | 303.083 | 5449.221 | 221.678 | 10.721 | 0.010 |
| | Diff (S1-S6) | | -1.631 | -2.360 | -8.076 | -0.108 | -0.002 |

[1] Genome Size (GS) estimates are in pg/2c. For MR19, GS estimates are from three S7 plants but sequence data were from three S6 plants.
[2] Values are estimates of Mb attributable to each genomic component, averaged across individuals within each landrace and generation.

**Table S9.** Estimates of the variance components and p-values with normalized read count data based on an ANOVA model and the B73 reference dataset. TEs were tested by location, where locations were split into overlapping genes (oTEs), withing 5kb of genes (gTEs) and non-genic (ngTEs).

| | Group | Landrace | Generation | Group X Gen | Race X Gen |
|---|---|---|---|---|---|
| gTE_v | 4.118 | 23.249 | 8.342 | 0.782 | 24.691 |
| gTE_p | 0.208 | 0.0643 | 0.0748 | 0.627 | 0.0578 |
| ngTE_v | 33.291 | 39.597 | 5.173 | 9.112 | 2.463 |
| ngTE_p | **9.70e-07** | **5.64e-06** | **8.87-03** | **1.44e-03** | 4.13e-01 |
| oTE_v | 23.076 | 25.995 | 9.989 | 12.385 | 9.937 |
| oTE_p | **0.000283** | **0.00265** | **0.00768** | **0.00364** | 0.0783 |

[1] The suffix v refers to the percentage of the total variance explained.

[2] The suffix p refers to probability. Bolded values are significant *p* = 0.05 after FDR correction for all tests in the Table.

**Table S10.** Estimates of the variance components and p-values with normalized read count data based on an ANOVA model and the W22 reference dataset. TEs were tested by location, where locations were split into overlapping genes (oTEs), within 5kb of genes (gTEs) and non-genic (ngTEs).

| | Group | Landrace | Generation | Group X Gen | Race X Gen |
|---|---|---|---|---|---|
| gTE_v | 14.201 | 71.603 | 2.664 | 5.600 | 1.128 |
| gTE_p | **4.73E-07** | **4.72E-11** | **3.93E-03** | **1.53E-04** | 4.21E-01 |
| ngTE_v | 15.085 | 68.953 | 3.448 | 5.982 | 1.149 |
| ngTE_p | **5.77E-07** | **1.03E-10** | **2.35E-03** | **1.79E-04** | 4.48E-01 |
| oTE_v | 13.018 | 65.258 | 5.568 | 8.121 | 1.925 |
| oTE_p | **3.83E-06** | **3.94E-10** | **4.77E-04** | **4.77E-04** | 2.74E-01 |

[1] The suffix v refers to the percentage of the total variance explained.

[2] The suffix p refers to probability. Bolded values are significant $p = 0.05$ after FDR correction for all tests in the Table.

**Table S11.** Estimates of the variance components and p-values with normalized read count data based on an ANOVA model and mapping to the B73 dataset. The model was tested separately for each of six different transposable element subfamilies (Helitrons, LINEs, SINEs, solo LTR elements, TIR and complete LTR elements).

| | Group | Landrace | Generation | Group X Gen | Race X Gen |
|---|---|---|---|---|---|
| Helitron_v | 30.648 | 39.440 | 5.492 | 7.971 | 4.984 |
| Helitron_p | **1.19e-06** | **5.83e-06** | **1.10e-02** | **0.00256** | 0.162 |
| LINE_v | 30.133 | 49.556 | 0.671 | 0.157 | 3.297 |
| LINE_p | **1.18e-05** | **1.18e-05** | 0.542 | 0.815 | 0.571 |
| SINE_v | 34.446 | 50.559 | 2.945 | 0.594 | 1.255 |
| SINE_p | **2.58e-07** | **4.26e-07** | 0.0427 | 0.440 | 0.815 |
| Solo_v | 0.319 | 70.660 | 4.828 | 9.938 | 2.142 |
| Solo_p | **0.622** | **2.06e-07** | 0.0174 | **0.00125** | 0.622 |
| TIR_v | 26.973 | 54.957 | 0.602 | 2.239 | 7.320 |
| TIR_p | **2.58e-07** | **6.45e-08** | 0.360 | **0.0427** | **0.0131** |
| LTR_v | 14.236 | 71.742 | 2.614 | 5.250 | 0.537 |
| LTR_p | **1.58e-06** | **4.30e-10** | 0.0152 | **0.000699** | 0.882 |

[1] The suffix v refers to the percentage of the total variance explained.

[2] The suffix p refers to probability. Bolded values are significant after FDR correction for all tests in the Table.

**Table S12.** ANOVA results based on counts to an independently annotated set of 22,530 full-length Sirevirus LTR retrotransposons. Hits were counted separately to the entire Sirevirus, the Internal region (Int), the LTR regions, and the ratio of hits to the LTR and internal regions. If unequal recombination were acting to remove LTRs, we expected that

| | Group | Landrace | Generation | GroupXGen | RaceXGen |
|---|---|---|---|---|---|
| Sirevirus_v[1] | 33.246 | 27.026 | 6.115 | 10.308 | 7.772 |
| Sirevirus_p[2] | **5.93e-06** | **5.83e-04** | **1.58e-02** | **2.63e-03** | 8.97e-02 |
| LTR_v | 47.515 | 50.548 | 0.258 | 0.632 | 0.146 |
| LTR_p | **9.420e-19** | **2.434e-17** | 3.674e-02 | **2.540e-03** | 6.782e-01 |
| Int_v | 29.724 | 23.321 | 7.114 | 11.866 | 9.812 |
| Int_p | **2.77e-05** | **0.00263** | **0.0158** | **0.00263** | 0.0753 |
| Ratio_v | 45.812 | 52.337 | 0.00679 | 0.00615 | 1.35 |
| Ratio_p | **6.61e-21** | **5.46E-20** | 0.733 | 0.733 | **2.77e-05** |

[1] The suffix v refers to the percentage of the total variance explained.

[2] The suffix p refers to probability. Bolded values are significant after FDR correction for all tests in the Table.

**Table S13:** Numbers of SNPs examined in each line for which the parental (P) generation was inferred to be heterozygous.

| Line | Generation and Individual numbers | Number of SNPs without missing data | Number of heterozygous SNPs in parent | Heterozygous SNPs in parent with SIFT information |
|---|---|---|---|---|
| **MR01**[1] | S1 3<br>S5 1<br>S6 2 | 584,334 | 64,000 | 2,397 |
| **MR08** | S1 3<br>S6 3 | 612,760 | 71,322 | 2,929 |
| **MR09** | S1 2<br>S6 3 | 5,555,006 | 891,494 | 31,721 |
| **MR18** | S1 3<br>S6 3 | 293,219 | 28,267 | 1,350 |
| **MR19** | S1 3<br>S6 3 | 294,243 | 28,438 | 1,648 |
| **MR22** | S1 3<br>S6 2 | 5,014,350 | 831,324 | 29,807 |

[1] For line MR01, the generation 5 individual was combined with the generation 6 individual for contrasts to generation 1 individuals.

**Table S14:** Estimates, Standard Errors, and Z- and P- values to test for differences between pairs of estimates of the proportion of derived alleles, as shown in Figure 3B. The tests are based on a GLM model (see Methods).

| Linear hypothesis on the derived allele proportion | Estimate | Pr(>\|z\|) |
|---|---|---|
| non coding S1 minus synonymous S1 == 0 | 0.008598 | 0.164 |
| non coding S1 minus non synonymous tolerated S1 == 0 | 0.062975 | <0.001 |
| non coding S1 minus non synonymous deleterious S1 == 0 | 0.310925 | <0.001 |
| non coding S1 minus non coding S6 == 0 | 0.058299 | <0.001 |
| synonymous S1 minus synonymous S6 == 0 | 0.037749 | <0.001 |
| non synonymous tolerated S1 minus non synonymous tolerated S6 == 0 | 0.095765 | <0.001 |
| non synonymous deleterious S1 minus non synonymous deleterious S6 == 0 | 0.189841 | <0.001 |

**Table S15:** Estimates, Standard Errors, and Z- and P- values to test for relationships between derived alleles and recombination, as illustrated in Figure 3B. The tests are based on a GLM model (see **Methods**).

| **Linear hypothesis on the derived allele proportion** | **Gen.** | **SNP type** | **Estimate** | **Pr(>\|z\|)** |
|---|---|---|---|---|
| high recombination minus low recombination == 0 | S1 | non synonymous deleterious | -0.029394 | 0.91757 |
| | | non synonymous tolerated | -0.001301 | 1.0000 |
| | | synonymous | 0.009305 | 0.99375 |
| | | non coding | 0.07036 | 1.29e-08 |
| | S6 | non synonymous deleterious | -0.134466 | 5.98e-06 |
| | | non synonymous tolerated | 0.045901 | 0.00595 |
| | | synonymous | -0.034818 | 0.05420 |
| | | non coding | 0.161881 | < 1e-10 |

**Table S16:** SNP coverage for sampled plants.

| Line | Individual | Depth (uniquely mapped) | Depth (all mapped) |
|---|---|---|---|
| MR01 | S1a | 2.24852 | 2.65529 |
| | S1b | 2.26967 | 2.6762 |
| | S1c | 2.05035 | 2.39219 |
| | S5a | 2.24052 | 2.60013 |
| | S6a | 2.2224 | 2.57922 |
| MR08 | S1a | 2.20168 | 2.58756 |
| | S1b | 2.169 | 2.53621 |
| | S1c | 2.19919 | 2.57436 |
| | S6a | 2.18397 | 2.5306 |
| | S6b | 2.24853 | 2.61444 |
| | S6c | 2.29712 | 2.67089 |
| MR09 | S1a | 3.50624 | 4.09764 |
| | S1b | 3.39456 | 3.96067 |
| | S6a | 3.62688 | 4.21669 |
| | S6b | 3.70907 | 4.32355 |
| MR18 | S1a | 2.01423 | 2.33699 |
| | S1b | 2.1866 | 2.56197 |
| | S1c | 2.10818 | 2.47707 |
| | S6a | 2.08716 | 2.40051 |
| | S6b | 1.97979 | 2.26576 |
| | S6c | 2.19901 | 2.54161 |
| MR19 | S1a | 2.30112 | 2.70889 |
| | S1b | 1.89324 | 2.18192 |
| | S1c | 2.1138 | 2.46023 |
| | S6a | 2.28322 | 2.6709 |
| | S6b | 2.15116 | 2.49936 |
| | S6c | 2.11943 | 2.45748 |
| MR22 | S1a | 3.16472 | 3.68149 |
| | S1b | 3.29707 | 3.84965 |
| | S1c | 3.53656 | 4.13367 |
| | S6a | 3.60864 | 4.20085 |
| | S6b | 3.46641 | 4.03131 |
| | S6c | 3.4293 | 3.97166 |

**Table S17:** Files used to used to define the five genomic components. Gene and TE entries link to gff files. For rDNA, Genbank numbers are listed as well as a link to files that define tDNAs. For Knobs and B-chromosomes, Genbank files are listed. See Methods that describe annotation of rDNA and knob components. Gffs of W22 and B73 genomes that were based on the result of rDNA and knob mappings are available on figshare.com.

| Component | File or Genbank Nos. |
|---|---|
| **Genes** | **B73:** ftp://ftp.ncbi.nlm.nih.gov/genomes/genbank/plant/Zea_mays/latest_assembly_versions/GCA_000005005.6_B73_RefGen_v4 **W22:** ftp://ftp.ncbi.nlm.nih.gov/genomes/genbank/plant/Zea_mays/latest_assembly_versions/GCA_001644905.2_Zm-W22-REFERENCE-NRGENE-2.0 |
| **TEs** | **B73:** http://ftp.gramene.org/CURRENT_RELEASE/gff3/zea_mays/repeat_annotation/ **W22:** ftp://ftp.ncbi.nlm.nih.gov/genomes/genbank/plant/Zea_mays/latest_assembly_versions/GCA_001644905.2_Zm-W22-REFERENCE-NRGENE-2.0 |
| **rDNA (plus tDNA)** | **rDNA:** M16267.1, U46616.1, AF013103.1, X03990.1, X01365.1, M10248.1, K01868.1, U46636.1, U46635.1, U46619.1 **tDNA:** http://lowelab.ucsc.edu/GtRNAdb/Zmays5/zeaMay5-tRNAs.fa |
| **Knobs (plus CentC repeats)** | AY530216.1, AY530287.1, AY530286.1, AY530284.1, AY530282.1, AY530280.1, AY530278.1, AY530276.1, AY530274.1, AY530272.1, AY530270.1, AY530268.1, AY530266.1, AY530264.1, AY530262.1, AY530260.1, AY530258.1, AY530256.1, AY530254.1, AY530252.1, AY530250.1, AY530248.1, AY530246.1, AY530244.1, AY530242.1, AY530240.1, AY530238.1, AY530236.1, AY530234.1, AY530232.1, AY530230.1, AY530228.1, AY530226.1, AY530224.1, AY530222.1, AY530220.1, AY530218.1, AY530285.1, AY530283.1, AY530281.1, AY530279.1, AY530277.1, AY530275.1, AY530273.1, AY530271.1, AY530269.1, AY530267.1, AY530265.1, AY530263.1, AY530261.1, AY530259.1, AY530257.1, AY530255.1, AY530253.1, AY530251.1, AY530249.1, AY530247.1, AY530245.1, AY530243.1, AY530241.1, AY530239.1, AY530237.1, AY530235.1, AY530233.1, AY530231.1, AY530229.1, AY530227.1, AY530225.1, AY530223.1, AY530221.1, AY530219.1, AY530217.1, DQ352544.1, AF071127.1, AF071125.1, AF071123.1, AF071121.1, AF071126.1, AF071124.1, AF071122.1, DQ186871.1, DQ186872.1, AF030939.1, AF030937.1, AF030935.1, AF030940.1, AF030938.1, AF030936.1, AF030934.1, M32533.1, M32534.1, M32532.1, M32530.1, M32528.1, M32526.1, M32524.1, M32522.1, M35408.1, M32531.1, M32529.1, M32527.1, M32525.1, M32523.1, M32521.1, AY173950.1 |
| **B Chromosome Repeats** | S67586.1, EF190064.1 |

**Table S18:** Results of mapping to a database consisting of only knob queries. The counts are not adjusted for overlapping hits with Transposable Elements, as with the genomic queries. The counts were normalized across libraries using BUSCO counts from either B73 or W22, with consistent results.

| | Group | Landrace | Generation | Group X Gen | Race X Gen |
|---|---|---|---|---|---|
| B73_v | 21.780 | 11.80 | 0.239 | 0.545 | 1.565 |
| B73_p | **2.43E-12** | **2.31E-08** | 1.74E-01 | **3.44E-02** | **1.58E-02** |
| W22_v | 14.448 | 8.339 | 0.132 | 0.402 | 0.393 |
| W22_p | **1.69E-11** | **7.86E-08** | 3.45E-01 | 5.34E-02 | 4.14E-01 |

[1] The suffix v refers to the percentage of the total variance explained.
[2] The suffix p refers to probability. Bolded values are significant at $p$ = 0.05 after FDR correction for the number of tests within each reference normalization.

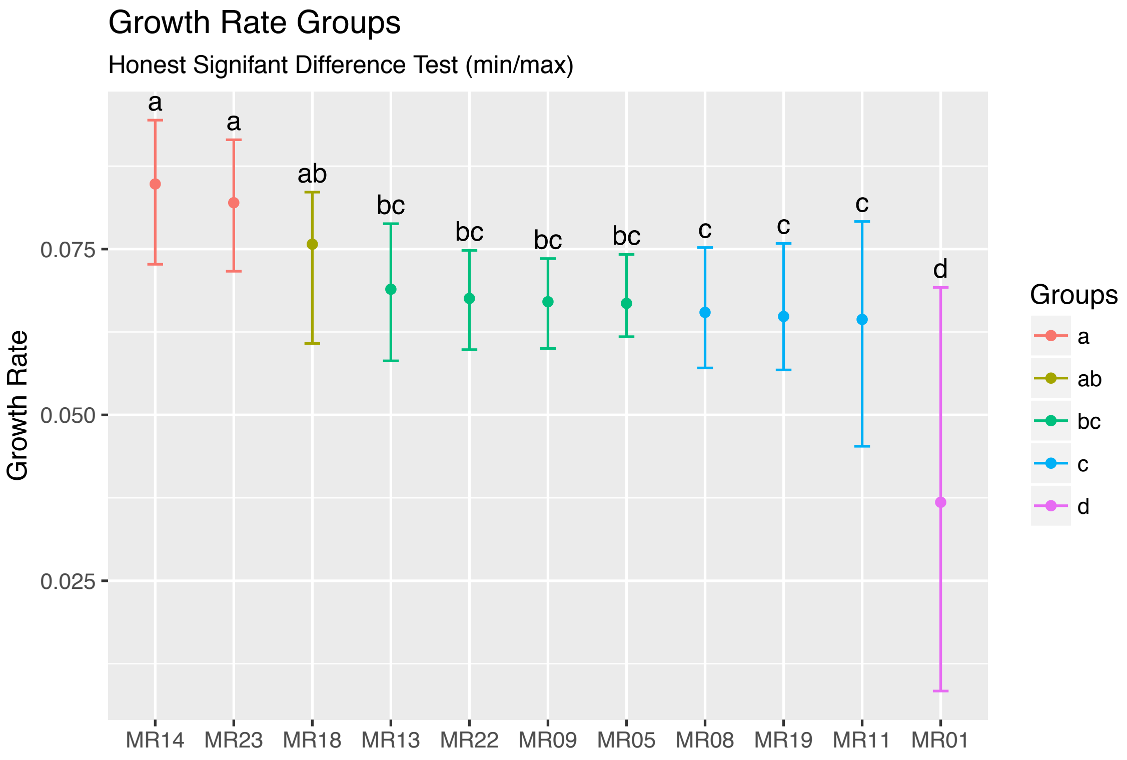

**Figure S1:** A comparison of growth rates among the 11 lines, based on all surviving plants from each at the end of 45 days. Each bar represents the estimated growth rate for each line, with the whiskers representing the minimum and maximum estimated growth rates for plants within the line. The letters above the bars denote statistical groups, based on the Honest Significant Difference Test.

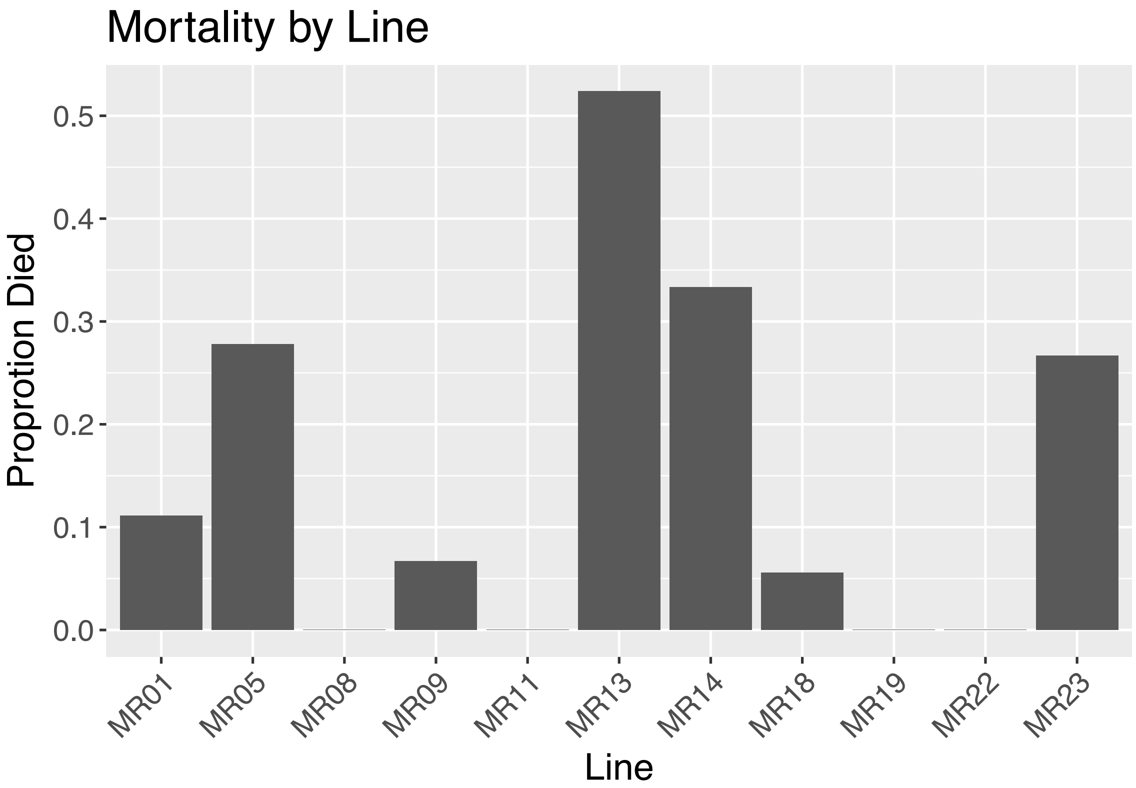

**Figure S2:** A report of mortality among the different lines. Mortality varied significantly among lines (binomial regression; $p < 0.001$).

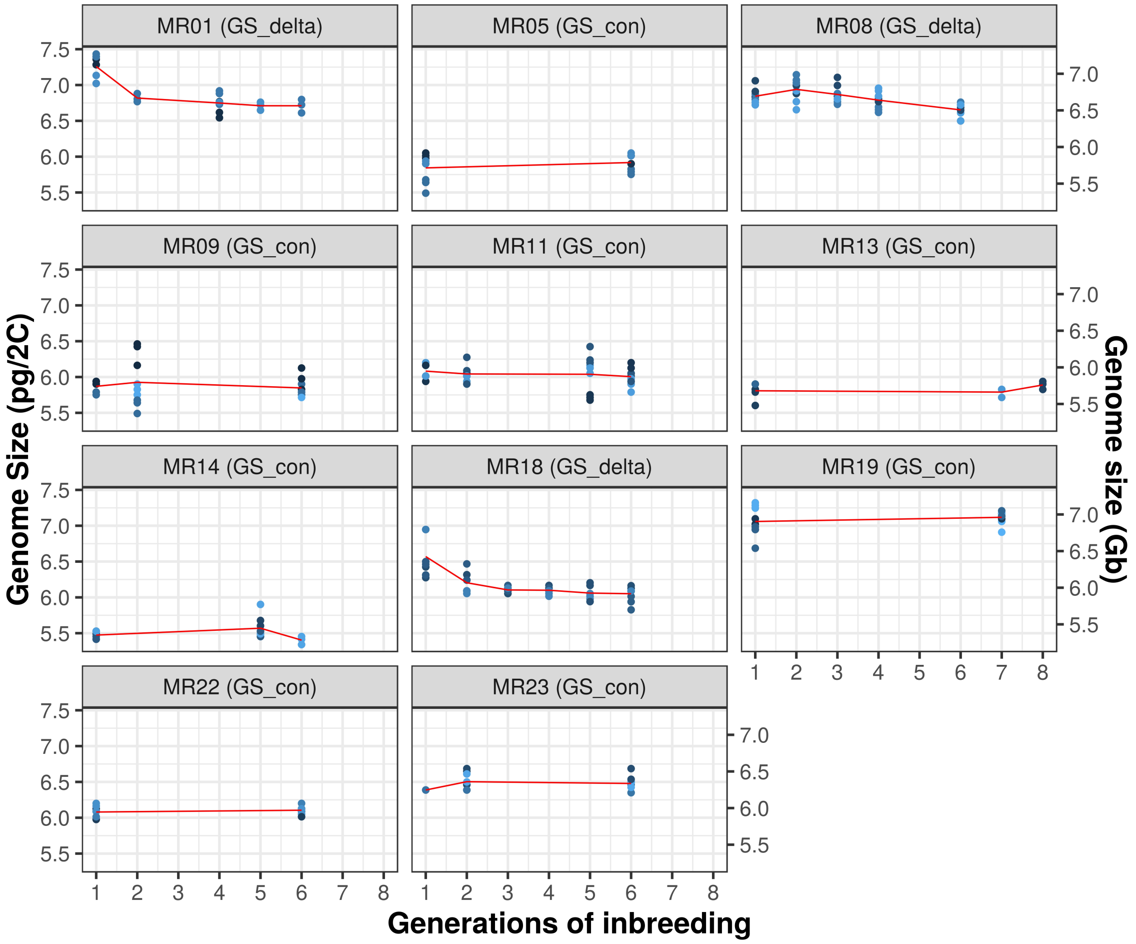

**Figure S3:** Flow cytometry data from Table S2 plotted per generation for each line separately. The conversion from pg/2C to Gigabases (Gb) is based on a conversion of 978 Mb per pg. GS_con refers to lines without a detectable change in genome size; GS_delta refers to lines with detectable losses of genome size.

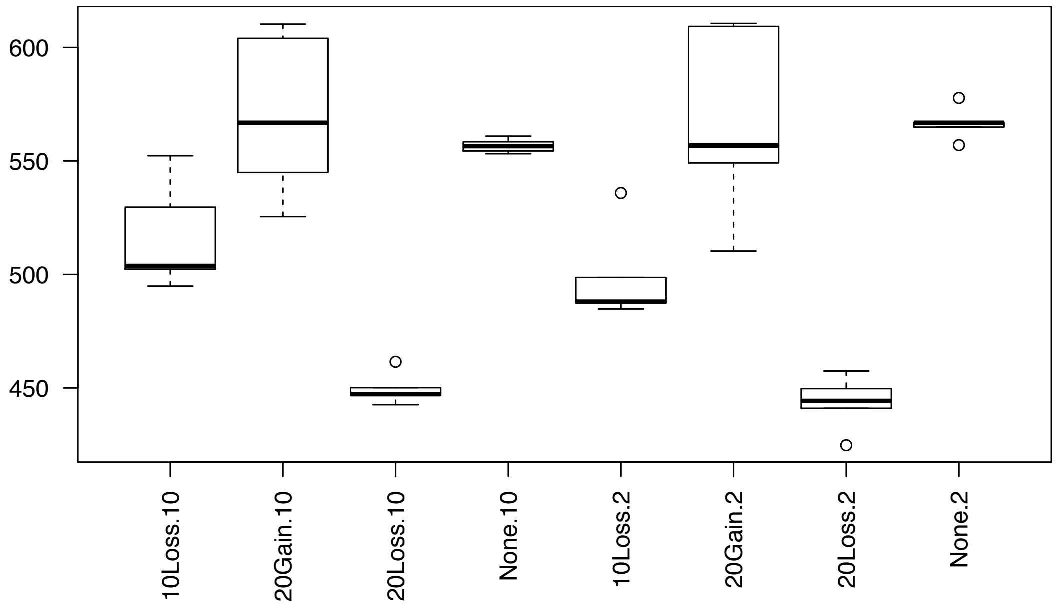

**Figure S4:** Simulation results for BUSCO method on chromosome 10, where either 10% or 20% of TEs were simulated as lost (10Loss, 20Loss), 10% of TEs were gained (10Gain) or there was no change to the genome (None). The simulations were run with either 10X or 2X coverage. The simulation parameters are indicated in the x-axis labels; for example, "10Loss.2" simulated the loss of 10% of TEs with genome coverage of 2x. The boxplots represent results from 1000 simulations. The black bar within the boxplot is the median, and the whiskers represent upper and lower quartiles.

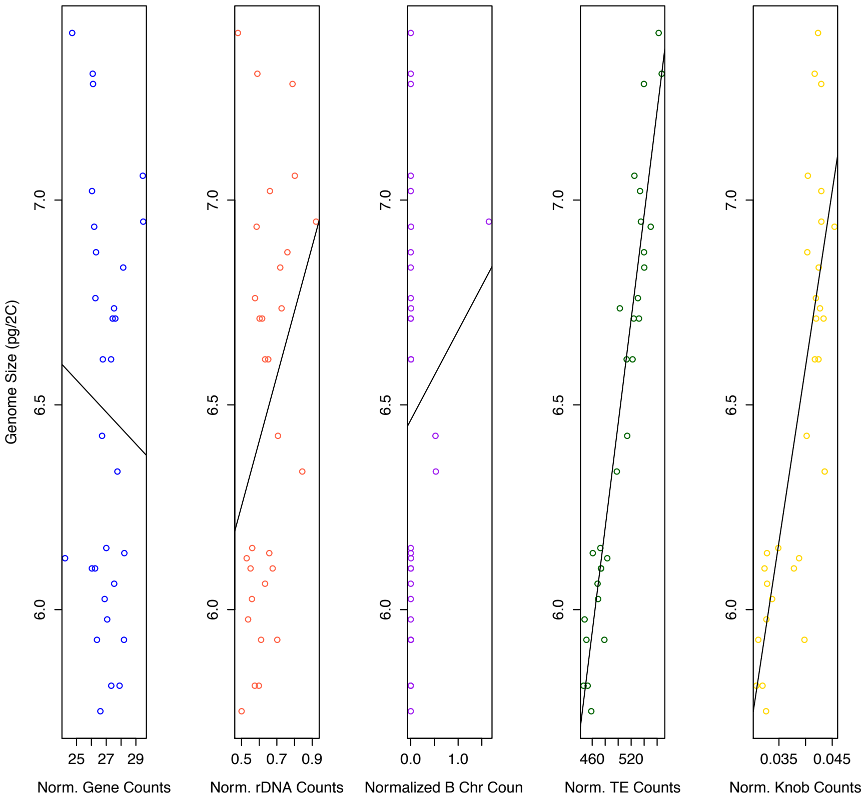

**Figure S5:** Trendlines for linear models comparing GS against normalized counts for each of five genomic components independently. The five components are all non-Busco genes, rDNA repeats, B-chromosome repeats, TE counts and knob repeats. Regressing each component separately, there was not a significant relationship to GS for genic content ($r^2$=-0.027, p=0.63), rDNA ($r^2$=0.079, $p$=0.07) or B-chromosome content ($r^2$=-0.015, $p$ = 0.45). There were strongly positive relationships between GS and both knob repeat content ($r^2$=0.662, $p$=4.5x$10^{-8}$) and TE content ($r^2$=0.901 p< $10^{-15}$). Reads were mapped to the B73 reference.

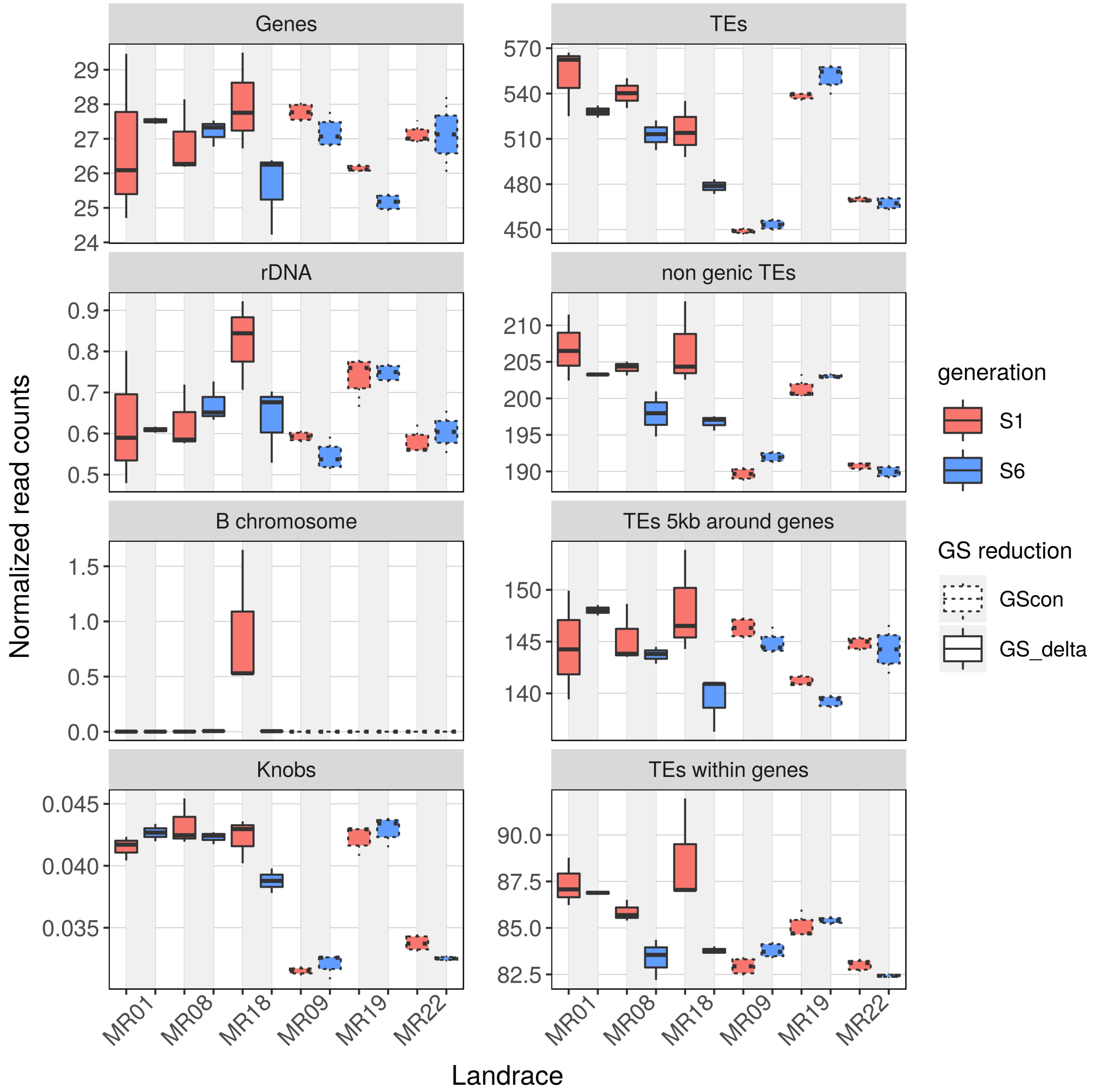

**Figure S6:** Plots of normalized read counts for various components of the genome for individuals from generations 1 and 6, based on mapping to the B73 genome. The boxes with solid outlines represent the GSΔ group that exhibited GS decreases, and the dotted outlines represent the lines with no change in GS (i.e., the $GS_{con}$ group).

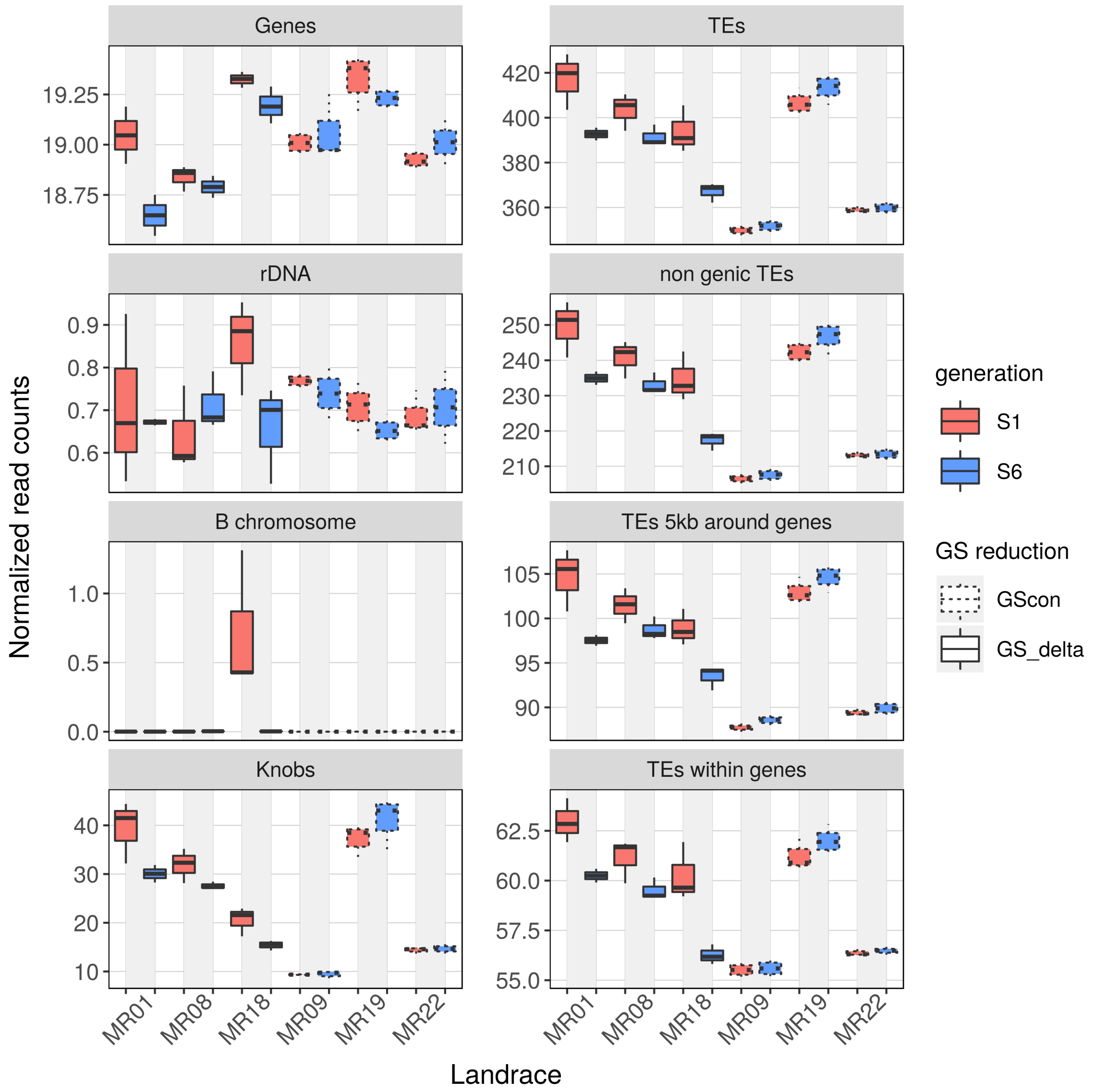

**Figure S7:** As per Figure S6, but for the W22 reference. Plots of normalized read counts for various components of the genome for individuals from generations 1 and 6. The boxes with solid outlines represent the GSΔ group that exhibited GS decreases, and the dotted outlines represent the lines with no change in GS (i.e., the $GS_{con}$ group).

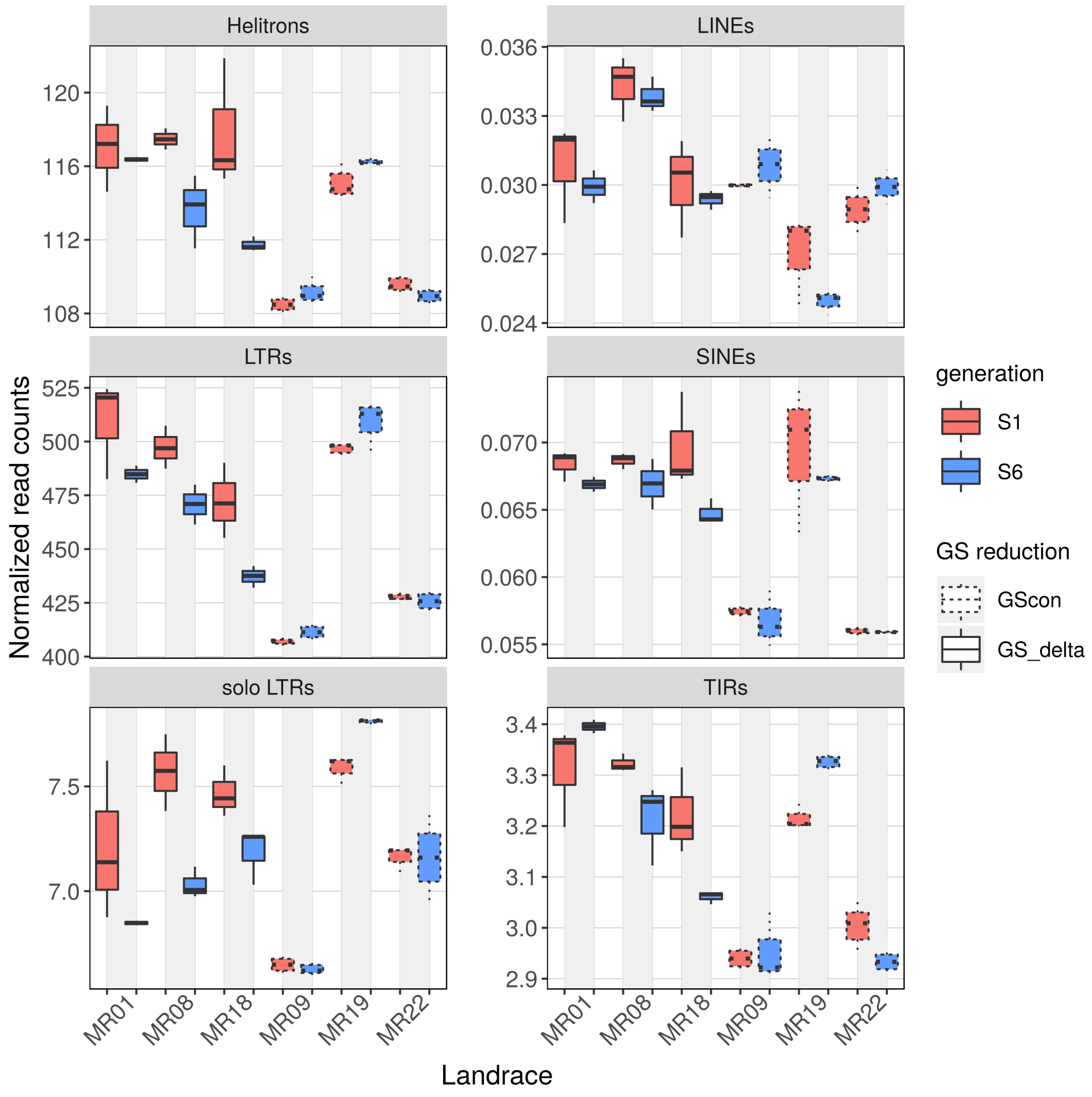

**Figure S8:** Plots of normalized read counts, based on mapping to B73, for various TE orders of the genome for individuals from generations 1 and 6. The boxes with solid outlines represent the GSΔ group that exhibited GS decreases, and the dotted outlines represent the lines with no change in GS (i.e., the $GS_{con}$ group). ANOVA analyses of these data are presented Iin Table S6.

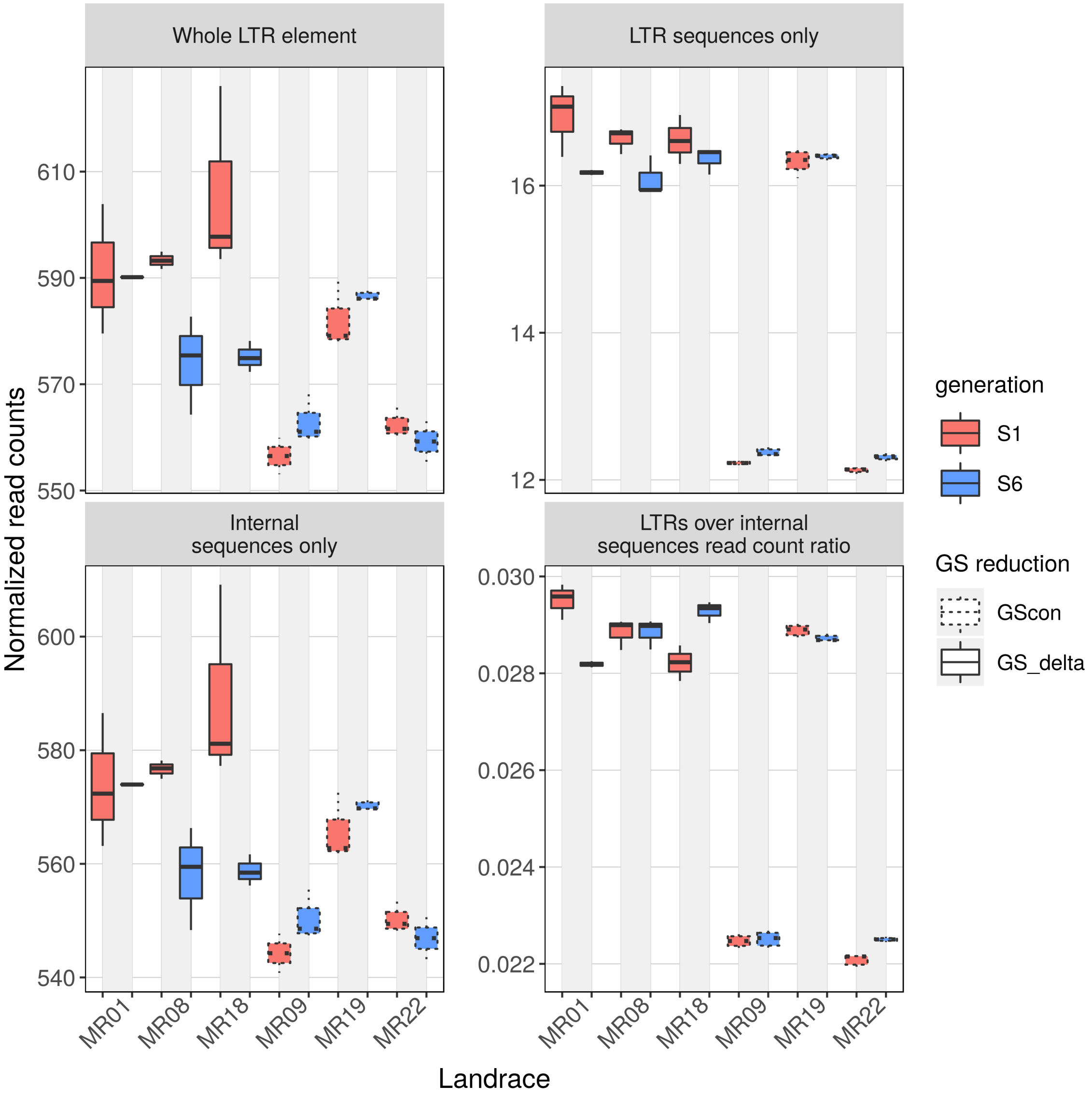

**Figure S9:** Plots of normalized read counts for 67,590 annotated Sireviruses. The boxes with solid outlines represent the GSΔ group that exhibited GS decreases, and the dotted outlines represent the lines with no change in GS (i.e., the $GS_{con}$ group). ANOVA analyses of these data are presented in Table S7.

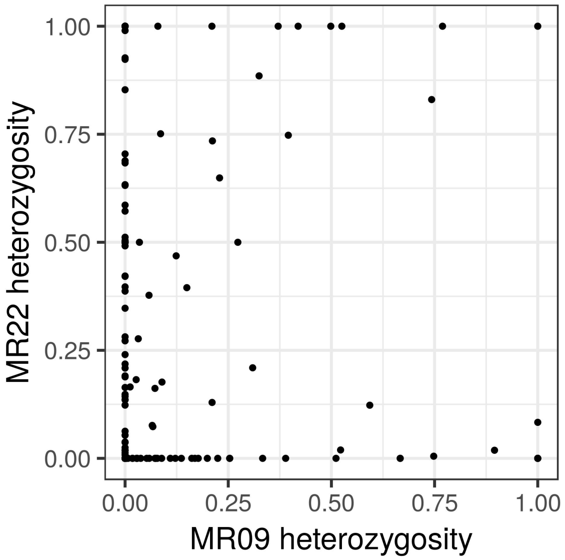

**Figure S10:** Comparing residual heterozygosity between the S6 generation of MR22 and MR09. Each dot represents a genomic region. See text for details of correlation statistics.

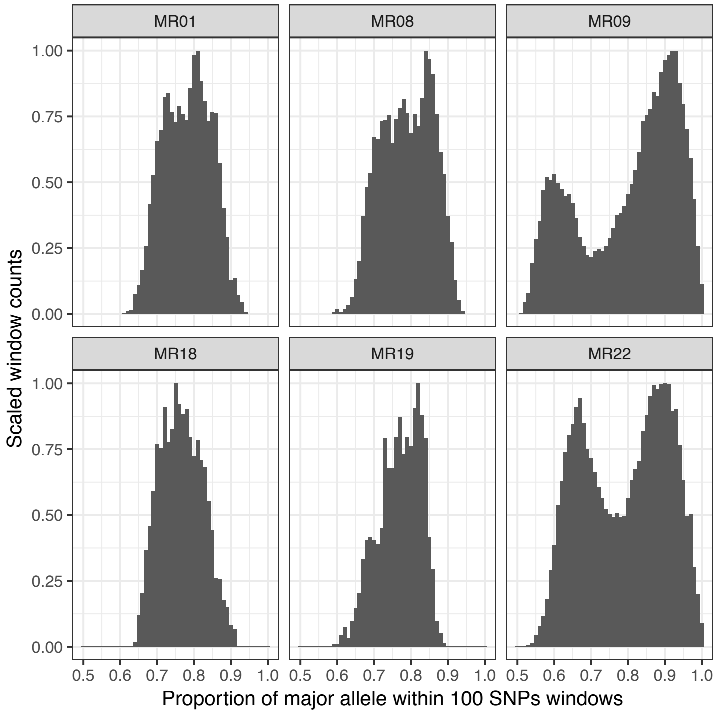

**Figure S11:** This should be the figure that shows the plots of windows vs. heterogyzity for all 6 lines… and indicates that only MR22 and MR09 have a bimodal distribution.

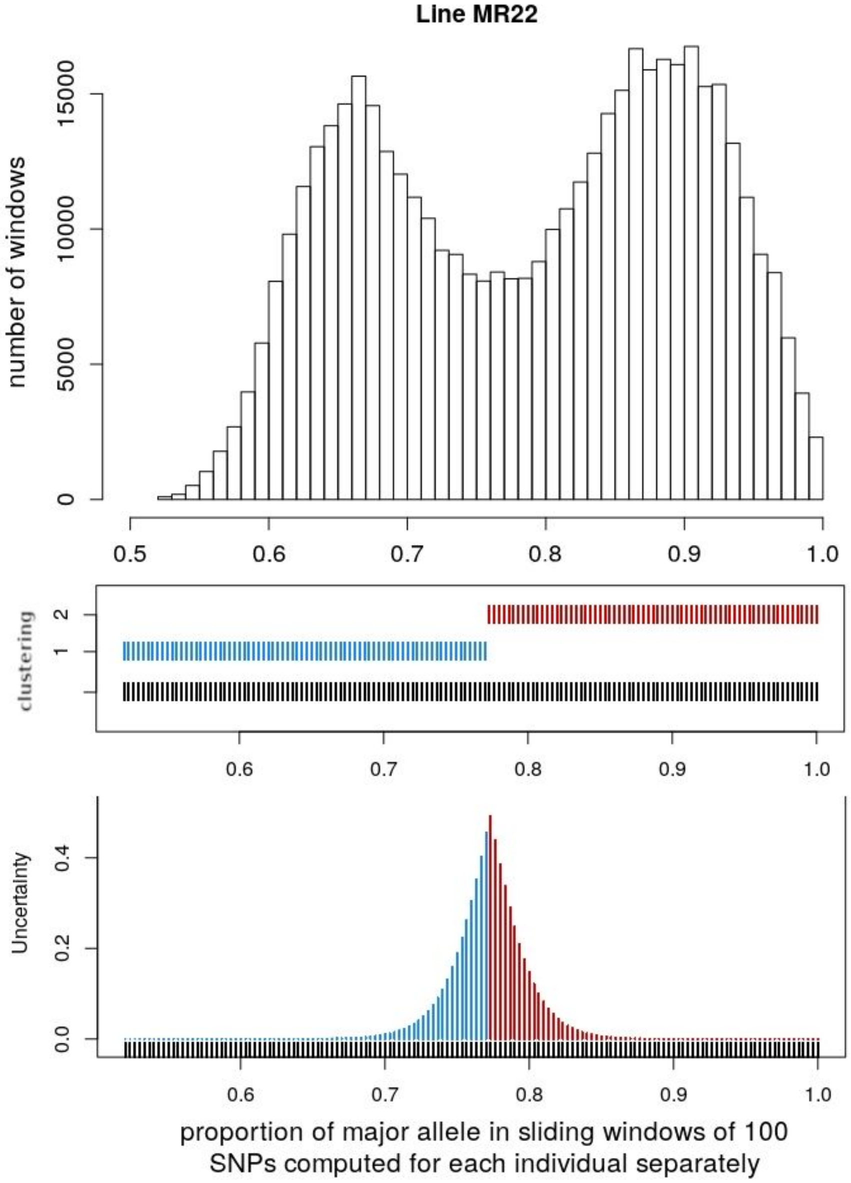

**Figure S12:** Properties of windows and heterozygous SNPs within the parent of MR22. The top graph shows that expected bimodal distribution of windows. The middle graph indicates results of Bayesian assignment. The bottom graph represents uncertainties in assignments.

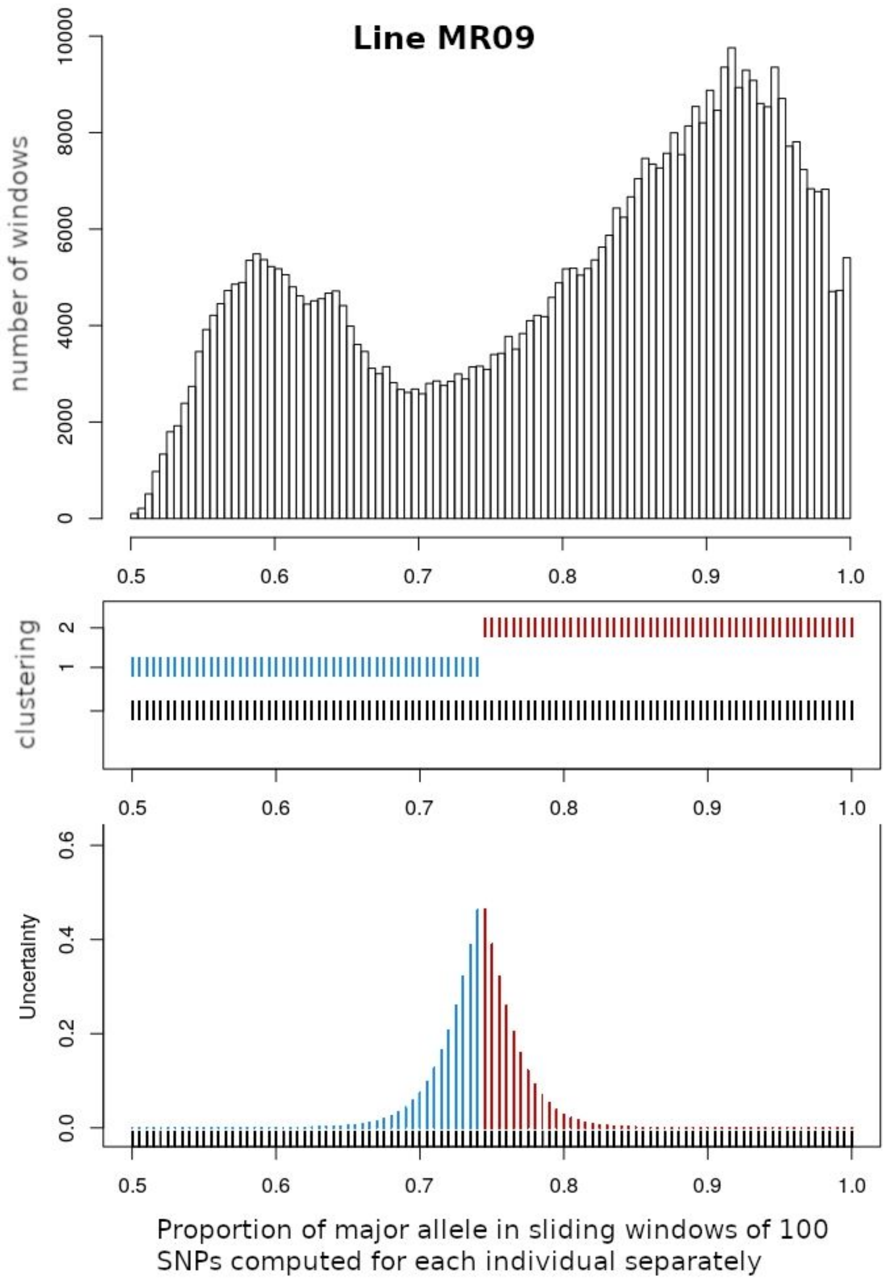

**Figure S13:** Properties of windows and heterozygous SNPs within the parent of MR09. The top graph shows that expected bimodal distribution of windows. The middle graph indicates results of Bayesian assignment. The bottom graph represents uncertainties in assignments.
